# The Separable Organization of Immune Transcriptional Responses

**DOI:** 10.64898/2026.09.09.750474

**Authors:** H. Jabran Zahid

## Abstract

Immune function depends on coordinated responses across diverse cell types, yet the organizing principles underlying this coordination remain uncertain. Here we analyze single-cell data of peripheral blood mononuclear cells from 12 donors exposed *in vitro* to 90 cytokines and find that, for a given donor and perturbation, transcriptional responses in one cell type can be transformed into corresponding responses in another through mappings that depend only on the identities of the two cell types. These cross-cell-type mappings admit a separable organization in which each donor–perturbation pair is represented by a response state shared across cell types, while donor- and perturbation-independent cell-type-specific response rules specify how that state is expressed. We formalize this organization as a linear shared-state model that captures most of the reproducible transcriptional response variance. This separation between shared response state and cell-type-specific response rules generalizes to unseen donors and perturbations and extends to longitudinal variation *in vivo*. Thus, our results reveal an organizing principle of immune coordination, with distinct transcriptional responses across cell types representing cell-type-specific expressions of a shared donor–level response state.

## Introduction

Biological organization is hierarchical, extending from molecules and cells to tissues and organs and ultimately to the level of the individual. Although experimental biology often studies these levels in isolation, organismal function and homeostasis depend on their coordination, motivating a systems-level understanding of biology [1, 2, 3, 4]. How molecular and cellular organization gives rise to organism-level biological state remains a fundamental challenge. Here we examine whether transcriptional change across cell types exhibits organization at the level of the individual.

The immune system provides a particularly tractable setting for studying the relationship between local cellular organization and the integrated biological state. Immune function is distributed across specialized cell types and tissues yet coordinated at the level of the individual [4, 5, 6]. The accessibility of peripheral blood has enabled matched molecular profiling of multiple immune cell types within the same donors [7, 8, 9, 10]. Recent studies have identified coordinated multicellular components of donor variation, including shared transcriptional programs, immune setpoints and longitudinal response structure [11, 12, 13, 14, 15, 16]. Building on these observations, we recently showed that transcriptional state across immune cell types is organized at the donor level, with this organization persisting over time and extending across tissues and molecular modalities [17]. Here we ask whether transcriptional responses are coordinated across peripheral immune cell types within donors and how this donor-level variation relates to cell-type-specific response patterns.

Changes in biological state provide a direct way to examine dynamic organization because they can reveal system structure that may not be apparent from static measurements alone [18, 19, 20]. Controlled *in vitro* stimulation, *in vivo* perturbations such as vaccination and longitudinal variation associated with aging have each been used to characterize changes across cell types, donors and molecular modalities [21, 22, 12, 13, 16]. *In vitro* perturbation is particularly informative because controlled genetic, cytokine and chemical interventions make causal effects and response structure more directly identifiable than in observational studies [18, 19, 23, 24, 25, 26]. Controlled perturbation studies have revealed substantial interindividual variation in immune responses, including effects of genetic variation on transcriptional responses that depend on stimulation context and cell-type-specific response differences [27, 28, 29, 30, 31]. We therefore use cytokine-stimulated peripheral blood mononuclear cells (PBMCs), which exhibit substantial donor-specific response heterogeneity [32], to examine the donor-level organization of transcriptional responses. We then test whether the same organization extends to natural longitudinal variation *in vivo* [16, 33].

Shared response organization across cell types may also be relevant to perturbation prediction, an important goal in computational biology and a focus of recent efforts to develop virtual-cell models [34]. Models have shown that responses can sometimes be predicted across perturbations, cell types and experimental contexts by learning transferable representations of response [35, 36, 37, 38, 39]. One potential source of such transferability is response variation associated with both perturbation and donor that is shared across cellular contexts but expressed differently in each population. Multi-view and latent-variable approaches provide a natural framework for identifying this structure by separating shared variation from view-specific components [40, 41, 42, 43, 44]. While these approaches can separate shared and context-specific variation, the organizing principles underlying donor-dependent perturbation responses across cell types remain unclear. Identifying such organization, if it exists, may provide a biologically structured basis for predicting responses.

Here we show that donor-dependent perturbation responses across peripheral immune cell types are not independently specified within each cell type, but instead exhibit a separable organization of donor–perturbation variation and cell-type-specific response structure. By separable organization, we mean that response variation can be separated into a component associated with donor and perturbation that is shared across cell types and donor- and perturbation-independent cell-type-specific rules that specify how that variation is expressed. We further show that the same organization extends to longitudinal transcriptional change *in vivo*.

## Results

We analyze a single-cell PBMC perturbation experiment comprising approximately 10 million cells from 12 donors exposed *in vitro* to 90 cytokines and phosphate-buffered saline (PBS) controls [32]. We focus on 11 major immune cell types that are broadly represented across donors and perturbations. We define each unique combination of donor, cytokine and cell type as a group. Within each group, cells are library-size normalized, log transformed and averaged to obtain a mean expression profile, retaining only groups containing at least 100 cells. We retain approximately 19,000 genes with nonzero mean expression in at least 50% of groups, preserving a broad transcriptional feature space without selecting genes based on variability or perturbation responsiveness.

For each stimulated group, we define the perturbation response as the difference between its mean expression profile and the corresponding donor- and cell-type-matched PBS mean profile. Each response vector therefore represents the group-level transcriptional change induced by a cytokine relative to the matched control. The eleven-cell-type dataset contains 10,892 observed donor–cytokine–cell-type response vectors, which constitute the primary units of analysis. Donor–cytokine combinations observed across multiple cell types provide matched cellular readouts of the same perturbation within the same donor, allowing us to compare response structure across cell types while preserving donor and perturbation correspondence. We analyze response vectors in a common principal-component representation that captures dominant variation while reducing measurement noise.

### Donor coordination of perturbation responses

We examine whether cytokine-induced responses are independently organized within each immune cell type or coordinated across cell types at the level of the donor. In particular, we demonstrate that cell types from the same donor exhibit related response patterns beyond the stereotyped effects shared across donors exposed to the same cytokine.

We apply principal-component analysis (PCA) jointly to all donor–cytokine–cell-type response vectors, defining a common transcriptional response basis across cell types, donors and perturbations. Although the responses span thousands of genes, a substantial fraction of total response variance is captured by the leading components of this basis (Fig. 1A). We retain the first 300 principal components as the representation for subsequent analyses. The fraction of reproducible response variation represented within this space is quantified separately below, and the results are insensitive to the exact number of retained components.

**Figure 1:**
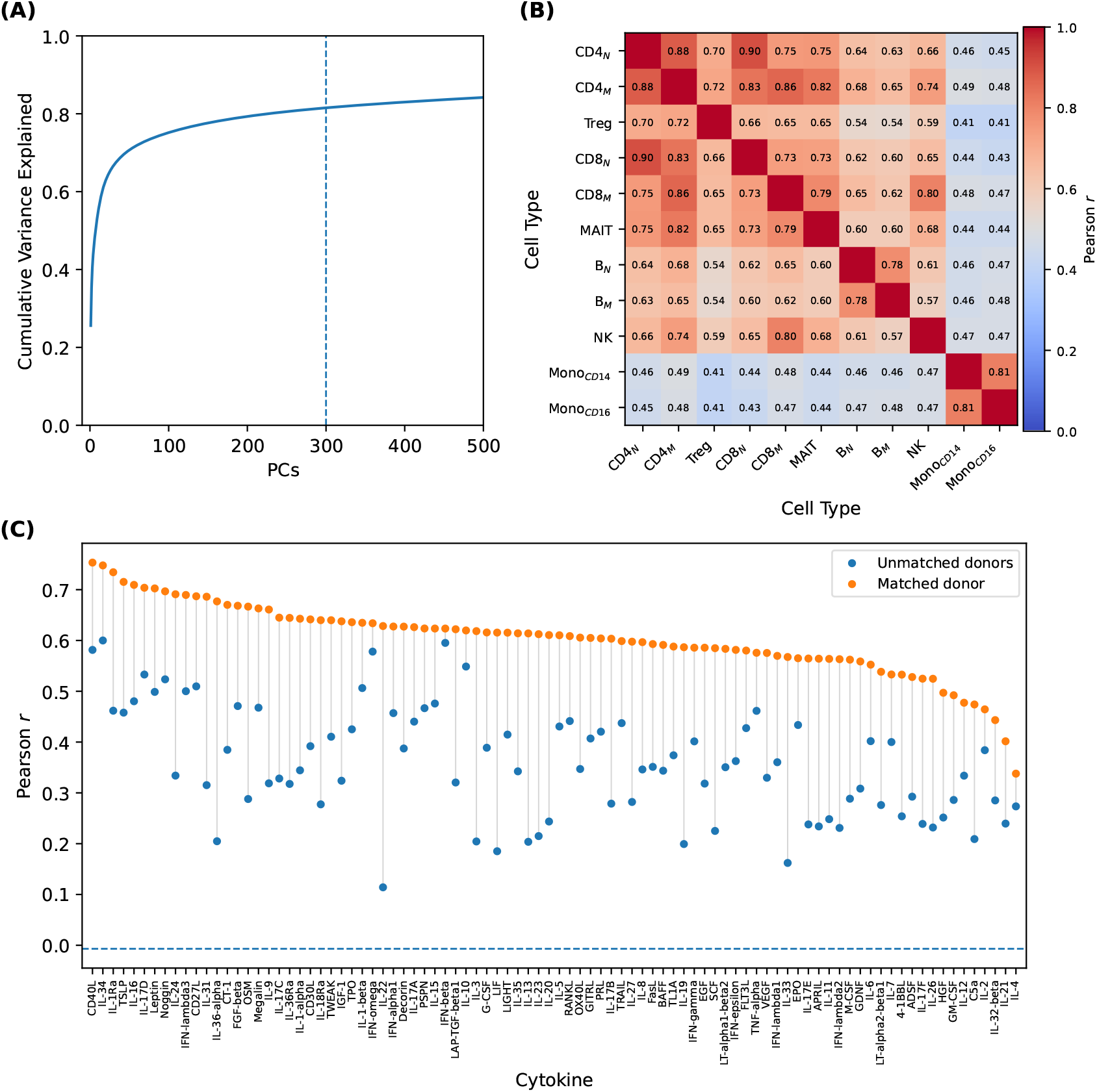
Cytokine-induced responses are coordinated across cell types within donors. **(A)** Cumulative variance explained by principal-component analysis of cytokine-induced response vectors across donors, cytokines and cell types. The dashed line marks the 300 principal components retained for subsequent analyses. **(B)** Median Pearson correlation between response vectors. Each matrix entry summarizes correlations across matched donor–cytokine cell type pairs. **(C)** Cross-cell-type response coordination for each cytokine. Matched donor (orange) values show the median pairwise cross-cell-type correlation within donors. Unmatched donor (blue) values show the corresponding correlation across donors exposed to the same cytokine. Correlations within matched donor are stronger for all 90 cytokines. The dashed horizontal line indicates the null median correlation obtained from responses for which both donor and cytokine identities are unmatched.

Within donors, response vectors show a reproducible pattern of similarity across cell types. For each matched donor–cytokine pair, we correlate the response vectors of two cell types and summarize these correlations across donor–cytokine observations (Fig. 1B). We find stronger correspondence among biologically related immune populations, including naive CD4 and CD8 T cells, memory CD4 and CD8 T cells, the two B-cell populations and the two monocyte populations, with weaker correspondence between more distinct lineages. This correspondence may reflect both a stereotyped response associated with a given cytokine and donor-specific variation coordinated across cell types. We distinguish these components by comparing cross-cell-type correlations within the same donor with correlations between different donors exposed to the same cytokine. The latter preserves the stereotyped cytokine response while removing donor correspondence. Across cytokines, cross-cell-type correlations are consistently higher within matched donors than between donors exposed to the same cytokine, while cross-cell-type responses are uncorrelated when both donor and cytokine identities are unmatched (Fig. 1C).

Together, these results show that cytokine responses contain a donor-dependent component coordinated across cell types beyond the stereotyped cytokine response shared across donors.

### Responses are related across cell types by stable mappings

Figure 1C shows that cross-cell-type response similarity contains both stereotyped cytokine structure and donor-dependent coordination. However, within donors, responses are not simply identical across cell types, motivating us to test whether their relationship may be better captured by stable cell-type-specific mappings.

For each source–target cell-type pair, we fit a linear mapping using responses matched by donor and perturbation and ask whether this mapping provides a stable description of the relationship between the two cell types. To test whether the learned mappings generalize beyond the donors and perturbations used to fit them, we use joint donor–perturbation cross-validation. Cytokines are divided into 10 folds, and within each perturbation fold each donor is held out in turn, yielding 120 training–test splits. Thus, donors and perturbations are held out simultaneously. For each split, the response principal-component basis and cross-cell-type mappings are learned using only the other 11 donors and 81 training cytokines. We then evaluate the learned mappings on the held-out donor–perturbation responses. Throughout, we refer to observed cell types used to generate a prediction as source cell types and the cell type whose response is being predicted as the target cell type.

We first examine the mapping between CD4 memory T-cell and CD14 monocyte responses, one of the more dissimilar cell-type pairs among the 11 populations. For each cross-validation split, we fit a linear ridge model using paired CD4 memory and CD14 monocyte response vectors from the training data. We then apply this mapping without modification to CD4 memory responses to predict CD14 monocyte responses from the held-out donor under the held-out cytokines. For each held-out donor–cytokine response, we quantify correspondence with the observed CD14 monocyte response in two ways: directly using the observed CD4 memory response and after transforming the CD4 memory response through the learned mapping. The learned mapping predicts the corresponding CD14 monocyte responses more accurately than direct, untransformed correspondence between the two cell types, showing that direct correspondence does not fully capture the stable relationship between their responses (Fig. 2A). Extending this analysis to all source–target pairs shows that learned mappings improve prediction over direct correspondence for every cell-type combination across held-out donor–perturbation states (Fig. 2B). Performance nevertheless varies by source and target, with T regulatory cells (Tregs) and monocytes generally showing lower prediction accuracy and a clear asymmetry in which they serve as stronger sources than targets.

**Figure 2:**
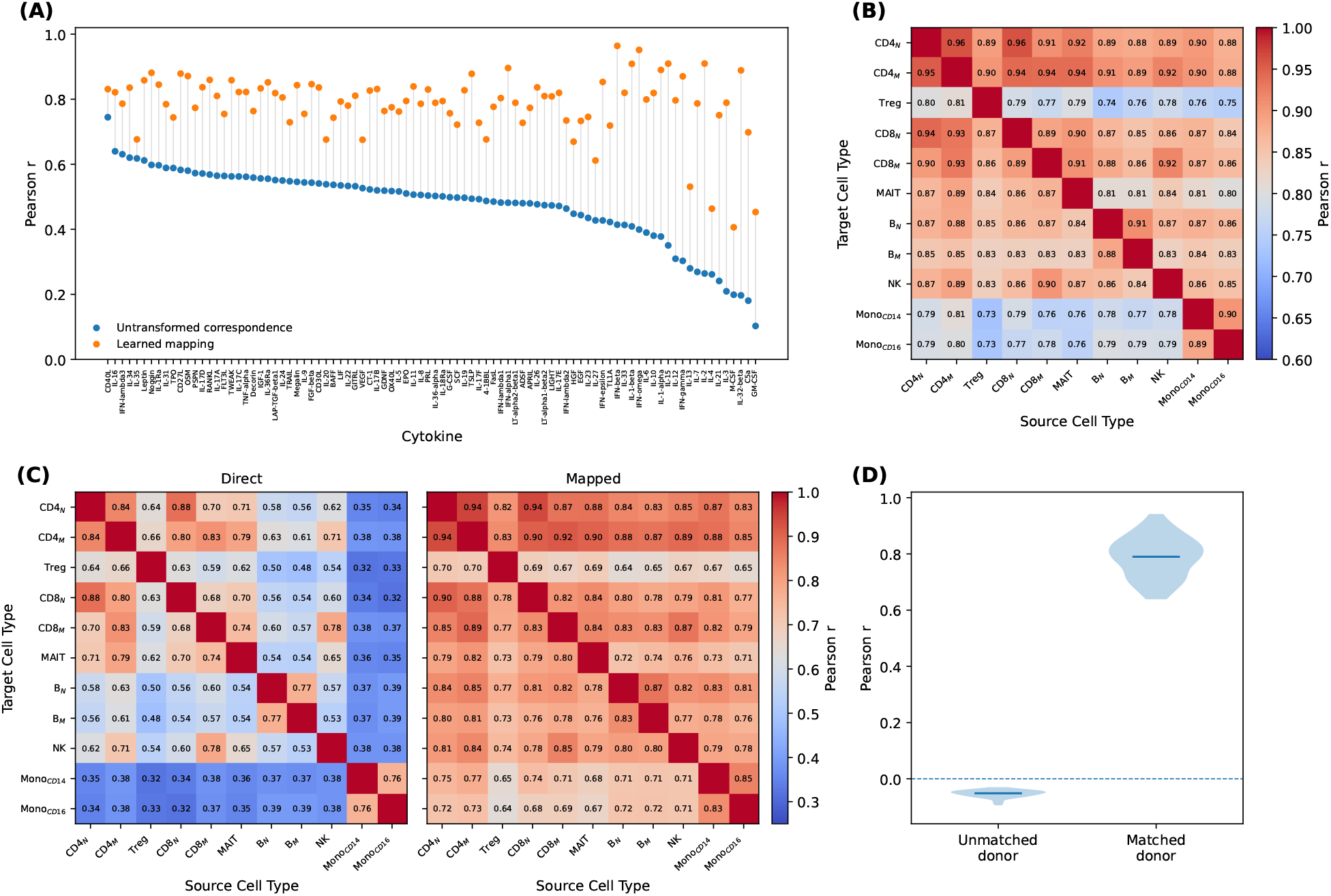
Responses in one cell type predict responses in others through linear cross-cell-type mappings that generalize across donors and perturbations. Throughout, each prediction is evaluated separately for a held-out donor–cytokine observation using the Pearson correlation across response PCs between the source-derived response and the corresponding target response. **(A)** Joint donor–perturbation prediction of CD14 monocyte responses from CD4 memory T-cell responses, comparing direct, untransformed correspondence (blue) with the learned mapping (orange). **(B)** Median prediction accuracy for cross-cell-type mappings learned and applied across all source–target pairs under joint donor–perturbation holdout. **(C)** Direct and mapped correspondence after subtracting the stereotyped cytokine response estimated from other donors exposed to the same held-out cytokine. The mapped deviations use the same mappings learned from the full training responses as in (B). **(D)** Mapped donor-specific response correspondence for matched versus unmatched donors exposed to the same cytokine. For each source–target mapping, correlations are summarized across held-out donor–cytokine observations; violins show the distributions of these summaries across the 110 source–target mappings.

The learned mappings could in principle be driven primarily by stereotyped cytokine responses shared across donors. We therefore ask whether the same mappings also capture the donor-specific component of response variation. To isolate this component, we subtract the stereotyped response to each cytokine. For each held-out donor, cytokine and cell type, stereotypy is estimated as the mean response of the other 11 donors exposed to that same held-out cytokine. These responses are used only to estimate stereotypy. The cross-cell-type mappings are not refit and are exactly those from the preceding analysis, learned from the other 11 donors under the 81 training cytokines. Thus, no responses to the held-out cytokine contribute to the mapping fit. After stereotypy is removed, direct correspondence between donor-specific response deviations is substantially weaker, while applying the same mappings learned from the full training responses markedly improves correspondence (Fig. 2C). Thus, the improvement from learned mappings is not confined to stereotyped cytokine responses. Rather, cell-type-specific mappings improve prediction of donor-specific deviations. Consistent with this interpretation, mapped donor-specific response deviations strongly correspond within matched donors but not across donors exposed to the same cytokine (Fig. 2D).

Figures 1 and 2 reveal three components of cross-cell-type response organization. First, different donors exposed to the same cytokine show correlated responses across cell types, reflecting a stereotyped component of cytokine response shared across individuals and cell types. Second, responses are more strongly correlated when both cytokine and donor are matched, revealing additional donor-dependent variation that is directly coordinated across cell types. Third, even within the same donor, responses are not identical across cell types; rather, learned cross-cell-type mappings substantially improve correlations over direct correspondence between the observed source and target responses, showing that donor-dependent response variation is expressed through non-trivial cell-type-specific mappings. The ability to predict target responses from all other source cell types indicates that donor-dependent response information is distributed across cellular populations rather than confined to any single cell type. At the same time, the generalization of these mappings supports the existence of donor- and perturbation-independent relationships between cellular responses. Together, these observations suggest a separation between variation associated with the donors and perturbations and the cell-type-specific rules through which that variation is expressed. In other words, for a given donor and perturbation, donor–perturbation information is shared across cell types, but each cell type expresses it differently. Importantly, these differences in cell-type-specific expression can be described by linear mappings that do not depend on the identity of the donor or perturbation. This motivates a model in which each donor–perturbation pair is represented by a response state shared across cell types, while donor- and perturbation-independent cell-type-specific response rules specify how that state is expressed.

### Separability of donor–perturbation response state and cell-type-specific response rules

The observations in Figure 2 motivate a separable model of transcriptional response organization. Figure 3A illustrates the inductive bias imposed by this model. Specifically, for each donor *d* and perturbation *p*, a single latent response state vector *Z*(*d, p*) is shared across all cell types, while each cell type *c* has a donor- and perturbation-independent linear decoder *W* (*c*) that maps the latent state into the observed response. We formalize this structure as a linear shared-state factorization,

**Figure 3:**
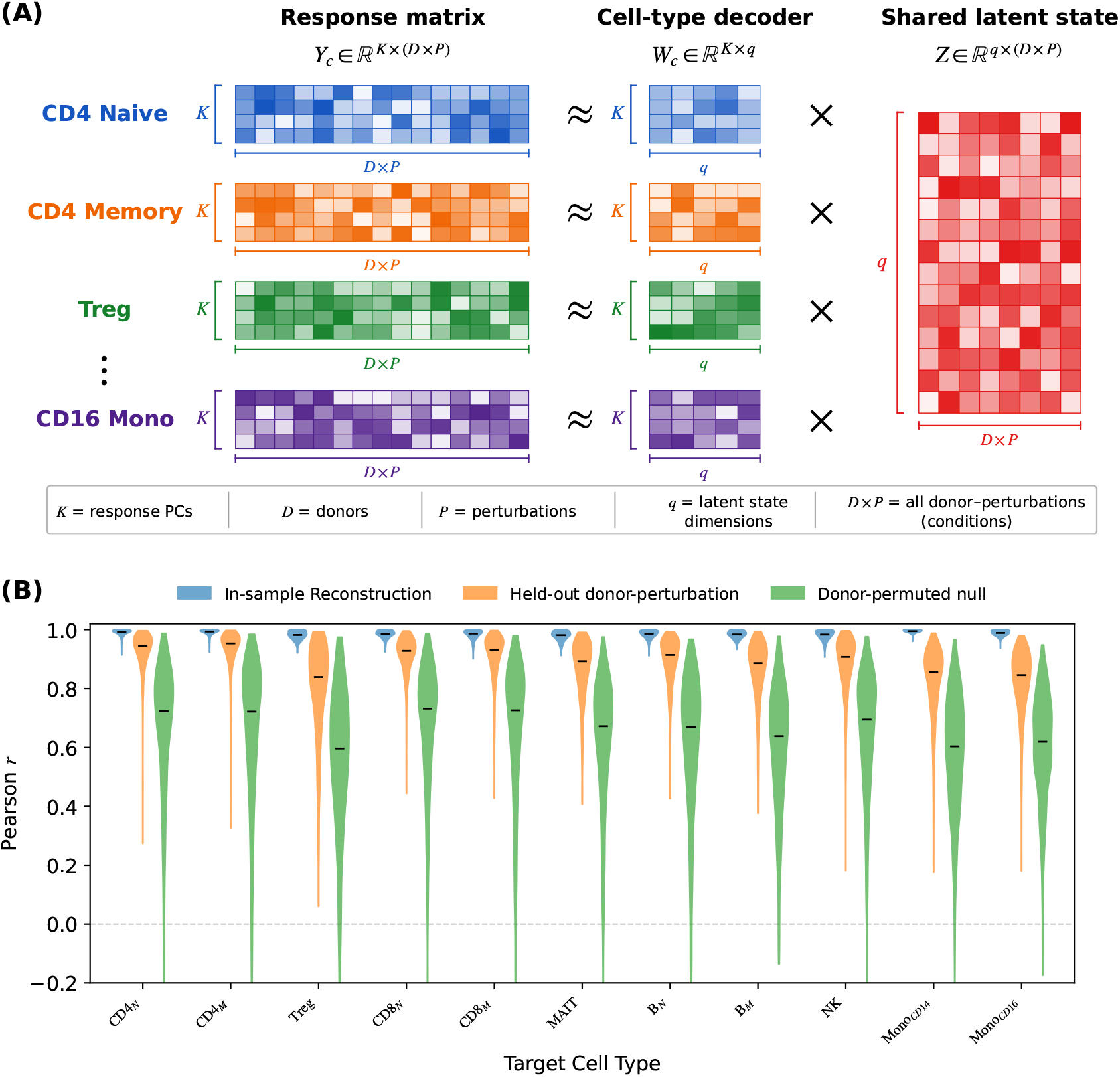
Perturbation responses separate into a shared donor–perturbation state and stable cell-type-specific decoders. **(A)** Schematic of the factorization *Y* (*c, d, p*) ≈ *W* (*c*)*Z*(*d, p*), in which a shared donor–perturbation state *Z*(*d, p*) is decoded into each cell-type response through *W* (*c*). **(B)** Reconstruction, joint donor–perturbation holdout prediction and donor-permuted null across target cell types. For prediction, each model is fit using the joint donor–perturbation cross-validation strategy described in the text. For each held-out donor– perturbation state, *Z* is inferred from all observed non-target cell types and decoded into the excluded target cell type using encoders and response rules learned without either the held-out donor or perturbation. For the null, donor correspondence is independently disrupted across cell types within each training cytokine before the model is refit and evaluated on the same correctly matched held-out donor–perturbation states. Violins show Pearson correlations between predicted and observed response vectors.

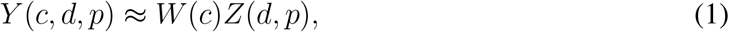

where *Y* (*c, d, p*) is the observed response of cell type *c* from donor *d* to perturbation *p*. The factorization therefore assigns donor- and perturbation-dependent variation to *Z*(*d, p*) and cell-type dependence to the decoder *W* (*c*). For each cell type, we additionally fit a linear encoder *E*(*c*) that maps the observed response *Y* (*c, d, p*) back into the latent-state space,

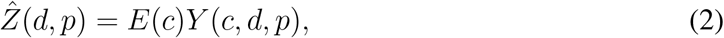

allowing the latent state to be inferred from observed cellular responses. *Z*(*d, p*), *W* (*c*) and *E*(*c*) are the key mathematical quantities of the learned model, with *Z* representing donor– perturbation response state and *W* and *E* providing the cell-type-specific decoding and encoding mappings, respectively. By construction, *Z*(*d, p*) is identical across cell types for a given donor–perturbation pair, while *W* (*c*) and *E*(*c*) are identical across donors and perturbations for a given cell type.

We test whether the separable model generalizes using the same joint donor–perturbation cross-validation strategy introduced above, in which donors and perturbations are simultaneously held out. Because *Z*(*d, p*) is a latent state rather than an independently observed quantity, the response state for a new donor–perturbation pair must be inferred from the observed cellular responses. For each held-out donor–perturbation pair, we therefore predict one target cell type, *c*_*t*_, at a time using only the responses of the other observed source cell types, *c*_*s*_. The inference procedure can be summarized as

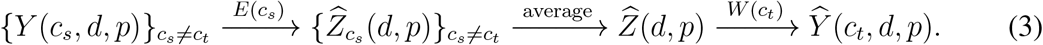

Specifically, each observed source response *Y* (*c*_*s*_, *d, p*) is mapped through its cell-type-specific encoder *E*(*c*_*s*_) to obtain a separate estimate 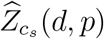 of the shared response state. These source-derived state estimates are then averaged across all observed non-target cell types to obtain a single inferred state 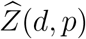. The inferred state is mapped through the target cell-type decoder *W* (*c*_*t*_) to predict the excluded target response 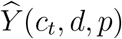. The target response *Y* (*c*_*t*_, *d, p*) is therefore never used to infer the state used to predict itself. In addition, the response PCA basis, source encoders and target decoder used for prediction are all learned only from the training donors and training perturbations, so none are fit using either the held-out donor or the held-out perturbation. This procedure therefore tests whether cell-type-specific encoders and response rules learned from training donors and perturbations generalize to unseen donor–perturbation states and support transfer from observed source cell types to an excluded target cell type. It also tests whether the shared response state of a new donor–perturbation pair can be inferred from the observed non-target cellular responses sufficiently well to predict the held-out target response.

The model accurately predicts responses across all target cell types under the joint donor– perturbation holdout (Fig. 3B). As a donor-correspondence null, we disrupt donor correspondence across cell types in the training data, refit the complete model and evaluate it on the same correctly matched held-out donor–perturbation states. The median Pearson correlation between predicted and observed response vectors is 0.91 and the model captures 83% of the variance within the retained 300-PC representation 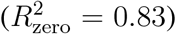, compared with a median correlation of 0.68 and 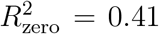 under the donor-correspondence null. The residual correspondence under the null largely reflects stereotyped perturbation responses shared across donors.

Together, these results show that the separable model generalizes to unseen donors and perturbations. The loss of predictive accuracy after disrupting donor correspondence further demonstrates that the shared state contains donor-dependent response information beyond stereotyped perturbation effects. Thus, the factorization captures the three components of response organization identified empirically above: stereotyped perturbation responses shared across donors, donor-dependent variation directly coordinated across cell types and donor-dependent variation expressed through non-trivial cell-type-specific mappings.

### The separable model captures most reproducible response variance

We next quantify how much reproducible transcriptional response variance is captured by the separable model. This provides a measure of the quantitative importance of the organization identified above and places the predictive performance in the context of the total biological signal present in the responses. Because observed response variance also contains measurement uncertainty, we use independent split-half estimates to distinguish reproducible response variation from measurement noise [45].

Across the total observed transcriptional response variance in gene space, 78% is reproducible across independent cell splits, while 22% is attributable to measurement uncertainty. We quantify how much of this response variance is captured by the model as a function of the PCA response dimensionality *K* (Fig. 4). Donor-cross-fitted PCA shows that at *K* = 300, the retained response space contains 95% of all reproducible gene-space response variance but only 10% of estimated total gene-space measurement uncertainty. Thus, within the PCA space, approximately 97% of the variance is estimated to be reproducible and 3% attributable to measurement uncertainty. Importantly, the separable model captures 86% of the reproducible variance within the first 300 response principal components, corresponding to an estimated 81% of all reproducible gene-space response variance.

**Figure 4:**
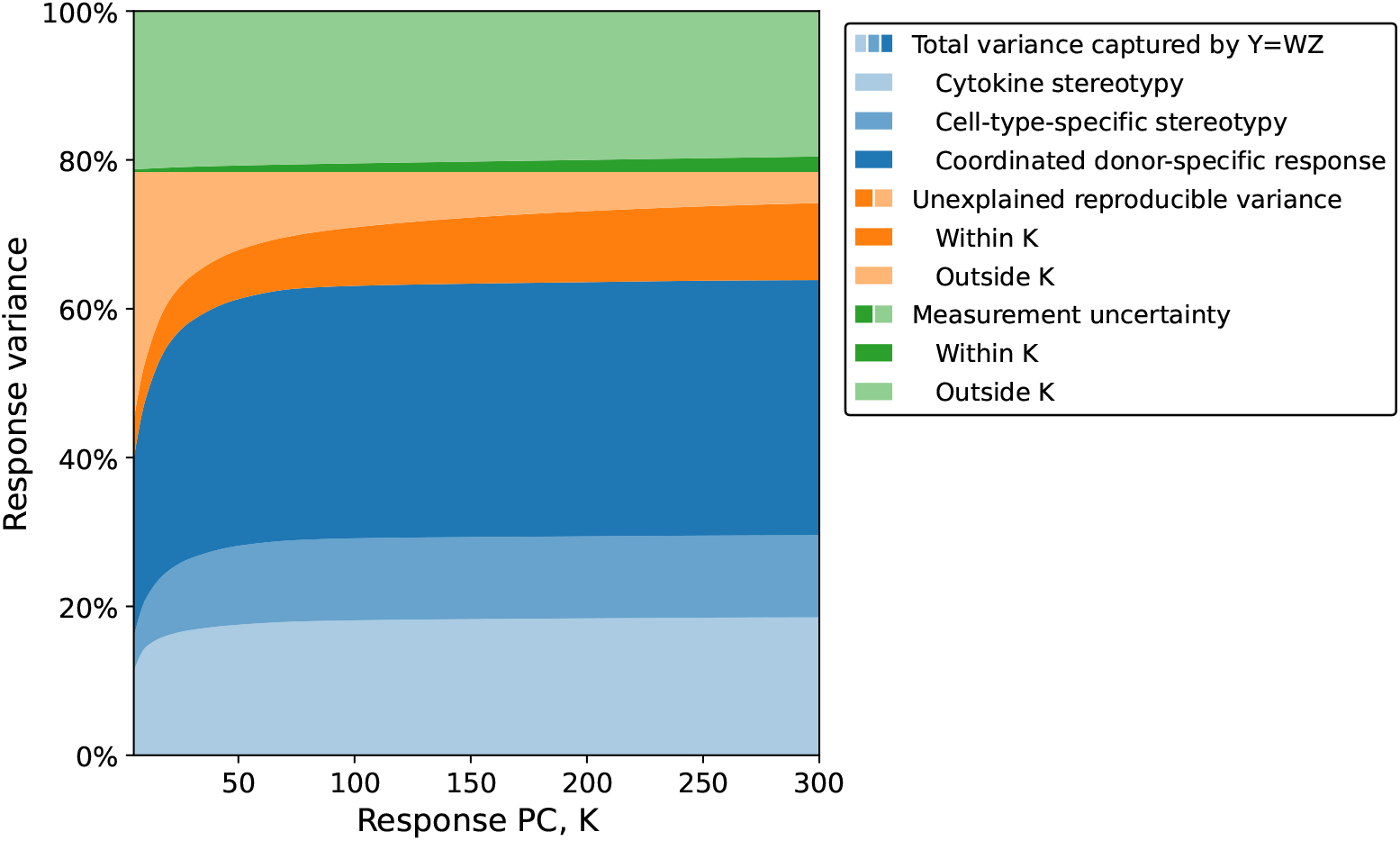
The separable model captures most reproducible transcriptional response variance. Stacked decomposition of total response variance as a function of retained principal components *K*. Blue regions partition variance captured by *Y* ≈ *WZ* into cytokine stereotypy, cell-type-specific stereotypy and coordinated donor-dependent response variation. Orange regions denote reproducible variance not captured by the model, while green regions denote measurement uncertainty. For both reproducible variance and measurement uncertainty, donor-cross-fitted PCA determines the fraction lying inside versus outside the PCA response space. At *K* = 300, the retained representation contains 95% of reproducible gene-space variance and our separable model captures 86% of this reproducible variance, corresponding to an estimated 81% of all reproducible response variance.

The variance captured by the separable model reflects the components of response organization identified above. At *K* = 300, stereotyped response structure accounts for 30% of total observed gene-space response variance. Of this, 19% reflects the mean response to each cytokine across donors and cell types, while an additional 11% reflects the cell-type-specific mean response to each cytokine across donors. Beyond these stereotyped components, the full separable model captures an additional 34% of total gene-space response variance associated with coordinated donor-dependent responses, making this contribution comparable in magnitude to the combined stereotyped response structure (Fig. 4). Thus, the separable organization accounts for most reproducible transcriptional response variation, with coordinated donor-dependent variation constituting a major component rather than a secondary effect.

### Distinct biological structure of separable organization

We next examine the biological structure represented by the shared response state and celltype-specific response rules. The factorization model assigns qualitatively different sources of variation to each. Donor- and perturbation-dependent variation is represented by a shared response state *Z*(*d, p*), whereas the mapping *W* (*c*) is cell-type-specific and specifies how that state is transcriptionally expressed. We therefore examine whether *Z* and *W* contain biological structure consistent with these distinct organizational roles.

We first examine the shared response states *Z*(*d, p*). To isolate perturbation-associated structure from donor-specific variation, we average *Z*(*d, p*) across donors for each cytokine. Similarity among these donor-averaged perturbation states reveals groups of cytokines with related response structure (Fig. 5A). This organization broadly recovers the cytokine groups defined by Oesinghaus et al. [32] (Fig. 5B), while also revealing broader hierarchical structure among perturbations. These independently defined cytokine groups form coherent regions within our representation, providing external biological context for *Z*. Perturbation identity therefore defines a major axis of organization within the shared response state. Conversely, after subtracting the mean state associated with each perturbation, residual states from the same donor are more similar across cytokines than states from different donors (Fig. 5C,D), indicating that *Z* retains donor response structure across perturbations. Thus, *Z*(*d, p*) contains both perturbation-associated structure and reproducible donor-specific response variation.

**Figure 5:**
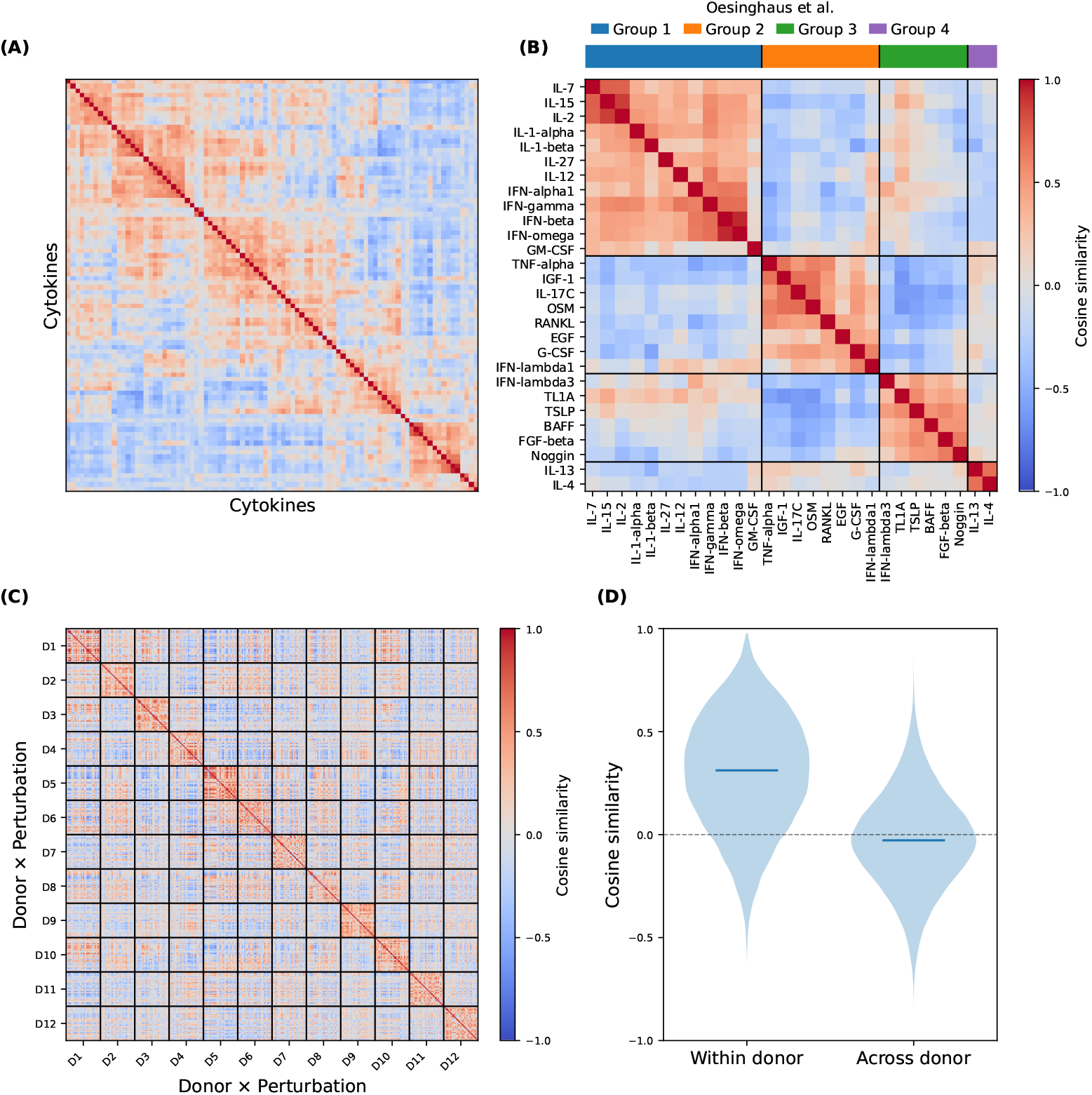
Shared response states encode perturbation and donor variation. **(A)** Pairwise cosine similarity among perturbation-level response states derived from *Z*(*d, p*), with cytokines ordered by hierarchical clustering of cosine similarity. **(B)** The subset of cytokines belonging to the four multi-cytokine groups defined independently by Oesinghaus et al. [32], ordered by group while preserving the *Z*-derived ordering within each group. **(C)** Pairwise cosine similarity among perturbation-centered donor response states. For each cytokine, the mean state across donors is subtracted from *Z*(*d, p*) to isolate donor-associated variation. Residual states are ordered by donor and then perturbation, with black lines separating donors. Similarity among residual states from the same donor indicates coordinated donor-specific response structure across perturbations. **(D)** Distribution of the pairwise cosine similarities shown in (C), separated into comparisons within the same donor and across different donors.

We next examine the cell-type-specific mappings *W* (*c*). Similarity is greatest among related lymphocyte populations, including the two B-cell populations, naive CD4 and CD8 T cells, memory CD4 and CD8 T cells and mucosal-associated invariant T cells (MAIT) while Tregs are more distinct. NK cells show greatest similarity to memory CD8 cells and the two monocytes are similar with one another and the most distinct from all other populations (Fig. 6A). Thus, the response rules are themselves biologically structured, with their relationships broadly recapitulating immune lineage organization. Together with the joint donor–perturbation generalization established above, this supports the interpretation of *W* (*c*) as a donor- and perturbation-independent, cell-type-specific response rule that maps a shared response state to the observed transcriptional response.

**Figure 6:**
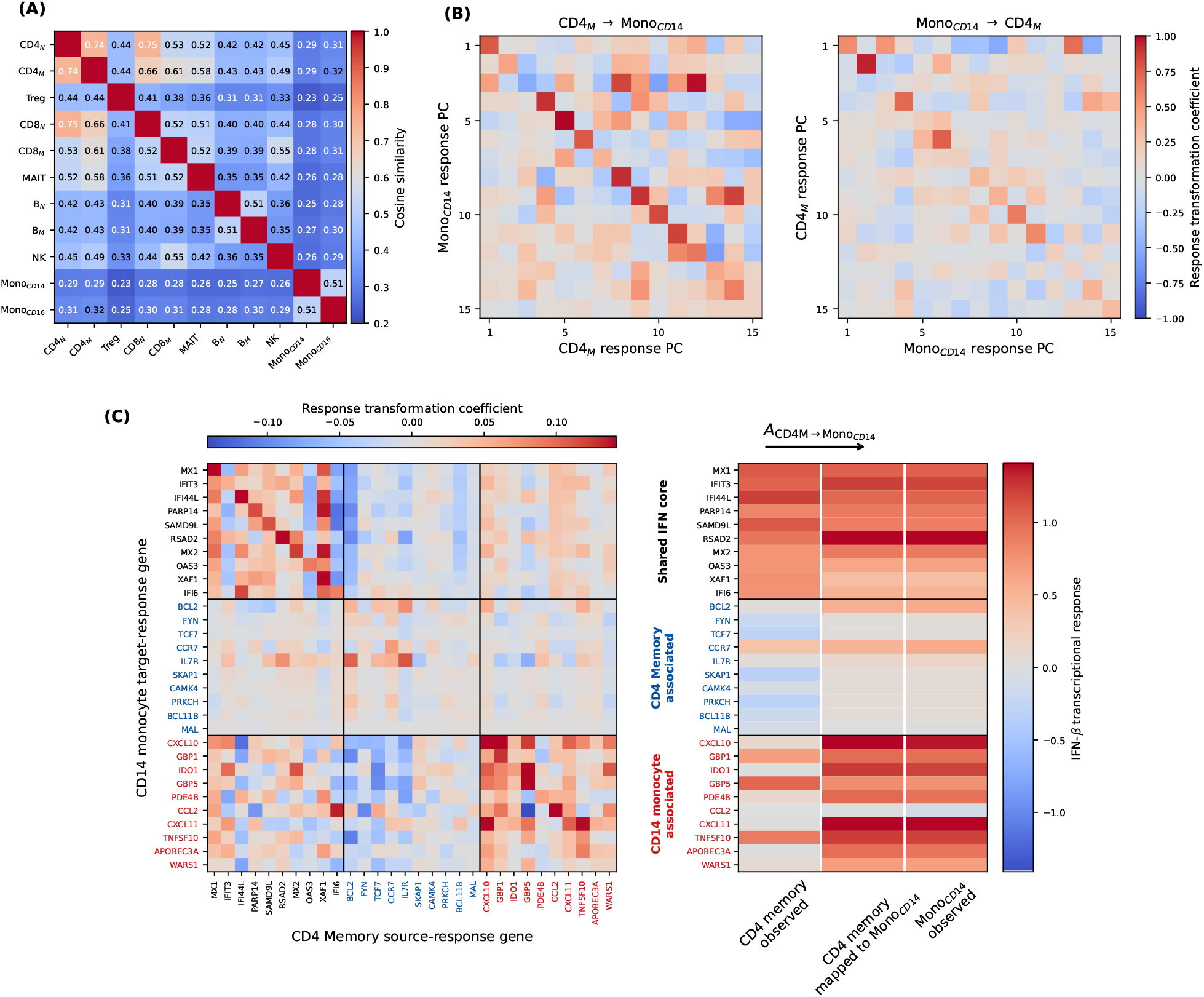
Cell-type-specific response rules map shared IFN-*β* response states to lineage-specific transcriptional programs. **(A)** Pairwise cosine similarity among fitted cell-type response rules *W* (*c*) reveals lineage-specific similarity. **(B)** Bidirectional mappings between CD4 memory T-cell and CD14 monocyte PCA response spaces, shown for the first 15 principal components, reveal structured cross-cell-type relationships. **(C)** Gene-space representation of the CD4 memory-to-CD14 monocyte mapping during IFN-*β* stimulation. The mapping separates shared IFN response genes from lineage-associated T-cell and CD14 monocyte inflammatory programs and maps the CD4 memory response state onto the corresponding CD14 monocyte transcriptional response.

To make the abstract mappings more concrete, we combine them into an operator that acts directly within the observed response PCA space. For a source cell type *c*_*s*_ and target cell type *c*_*t*_, the mapping proceeds as

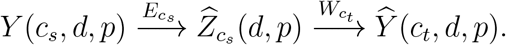

We therefore define the source-to-target operator as

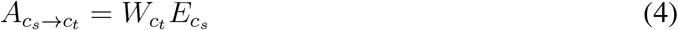

such that

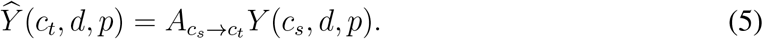

Thus, 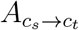 provides a direct mapping between cell-type responses in the observed response PCA space, directly describing how gene-space response patterns are transformed between cell types. Unlike the individual representations of *W, E* and *Z*, which depend on the latent-state coordinate system, *A* does not. Because both *W* and *E* depend only on cell type, the composed operator is also donor- and perturbation-independent. To visualize the structure of these mappings, we examine the CD4 memory-to-CD14 monocyte and CD14 monocyte-to-CD4 memory operators. These operators contain both diagonal and off-diagonal components and differ substantially by source-to-target direction (Fig. 6B), showing that cross-cell correspondence involves non-trivial, direction-specific mappings of response patterns. The asymmetric source– target generalization observed above (Figure 2B) and reported in other studies [46, 47] may reflect the direction-dependent structure captured by these operators.

To illustrate how these PCA response-space mappings relate to familiar transcriptional programs, we examine the CD4 memory-to-CD14 monocyte operator by projecting it from response-PC space back into gene space using a manually curated set of illustrative genes representing shared interferon response and lineage-associated CD4 memory and CD14 monocyte programs (Fig. 6C). The resulting operator contains recognizable block structure showing that CD4 memory-associated genes are broadly attenuated when mapping CD4 memory responses to CD14 monocyte responses, forming a near-zero band, whereas the shared interferon core and CD14 monocyte-associated genes have strong positive weights. Applied to the IFN-*β* response, the same operator maps the transcriptional pattern observed in CD4 memory T cells to the distinct pattern observed in CD14 monocytes. We emphasize that individual coefficients should not be interpreted as gene-to-gene regulatory interactions; rather, the operator describes how patterns of transcriptional changes in one lineage correspond to patterns of changes in another.

Together, Figures 5 and 6 show that the two components of the separable model carry distinct biological structure. *Z*(*d, p*) captures variation associated with the donor and perturbation, while *W* (*c*) reflects immune lineage relationships and specifies how a shared response state maps to transcriptional response.

### Shared-state organization extends to longitudinal immune variation

If the separable organization identified *in vitro* reflects a general principle of coordinated immune change, then naturally occurring longitudinal changes may exhibit a similar shared-state organization across cell types. We test this prediction in the independent Sound Life longitudinal PBMC cohort [16], which profiles 95 donors across nine scheduled visits with matched transcriptional measurements across nine immune cell types. For each donor, we define a cell-type-specific longitudinal trajectory for each pair of visits as the transcriptional change between those visits. We then define the donor-specific trajectory for that donor and visit pair as the collection of matched cell-type-specific longitudinal trajectories across observed cell types. Each donor therefore contributes multiple donor-specific trajectories corresponding to different visit pairs. As in the *in vitro* analysis, we represent these longitudinal changes in a common principal-component basis which is derived from all available cell-type-specific longitudinal trajectories across donors and visit pairs. We use the first 300 components to represent the response space. We then ask whether within-donor longitudinal change exhibits the same separable organization observed in *in vitro* perturbations.

We first examine longitudinal changes directly. Pairwise longitudinal differences show broad positive correspondence across the nine immune cell types within a given donor-specific trajectory. Consistent with the *in vitro* analysis, the strength of coordination varies among cell-type pairs and is weakest for comparisons involving monocytes (Fig. 7A).

**Figure 7:**
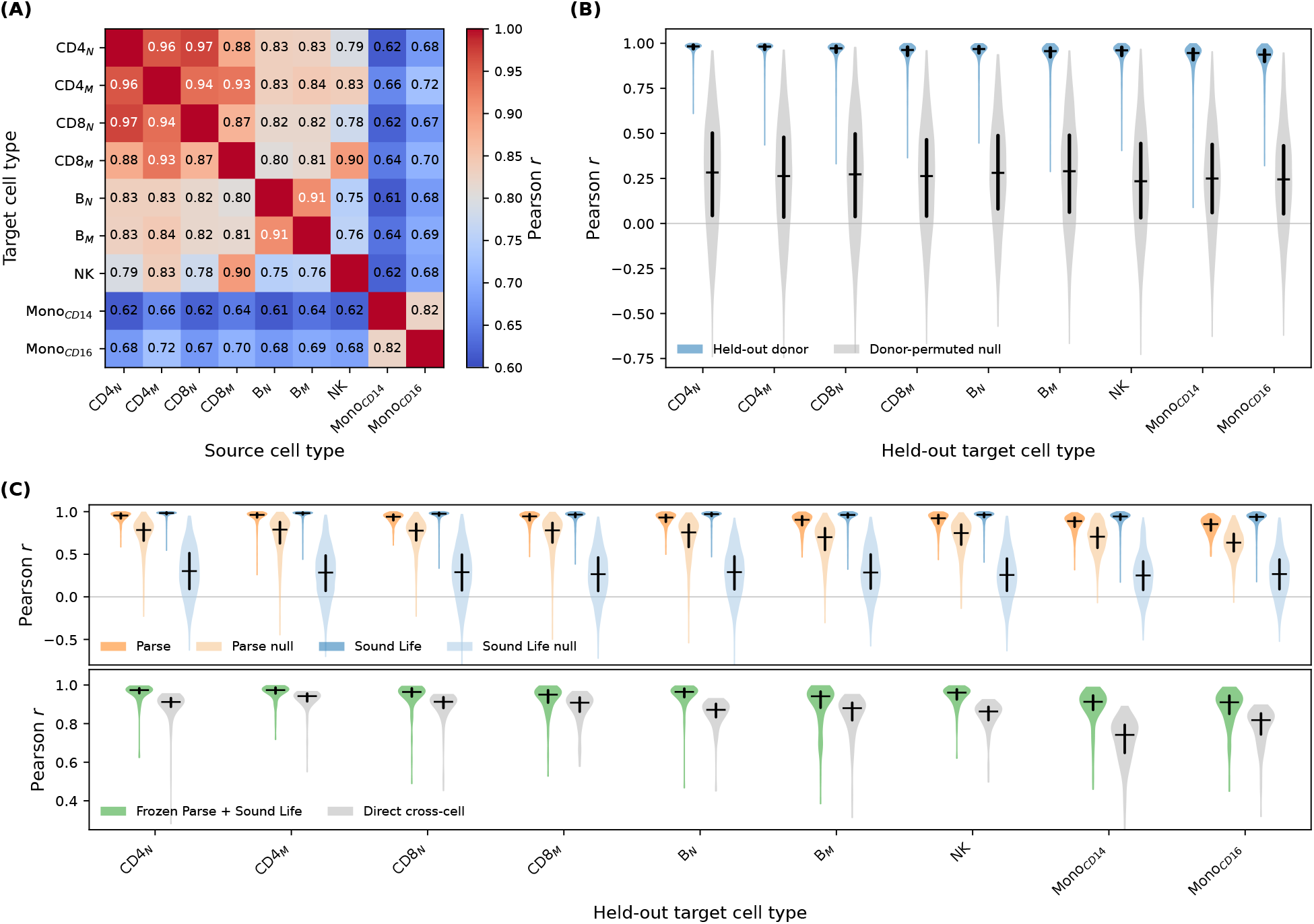
Separable organization extends to longitudinal immune change and generalizes across cohorts. **(A)** Direct correspondence of longitudinal changes across cell types within donor-specific Sound Life trajectories. Matrix entries show median Pearson correlations. **(B)** Prediction of longitudinal change in held-out Sound Life donors using the separable model. For each donor-specific trajectory and target cell type, the prediction is generated as before (Eq. 3). Violins show Pearson correlations between predicted and observed responses; the donor-permuted control disrupts donor correspondence across cell types within each training visit pair before model fitting. **(C)** Generalization across response settings. Top, a single separable model jointly fit to Parse cytokine responses and Sound Life longitudinal changes predicts excluded target responses in each cohort, with corresponding donor-permuted controls. Bottom, the jointly fit Parse–Sound Life model is frozen and applied without refitting to longitudinal rheumatoid arthritis data, where predictions are compared with direct cross-cell correspondence. Violins show Pearson correlations between predicted and observed responses.

We next test whether longitudinal immune change in Sound Life is described by the same separable factorization used for the cytokine perturbation data (Eq. 1). In strict five-fold donor-level cross-validation, all learned quantities, including the response PCA basis, shared-state model *Z*, cell-type response rules *W* and encoders *E*, are fit using training donor-specific trajectories only. As before, for each donor-specific trajectory and target cell type from a held-out donor, responses from the other observed cell types within that same trajectory are used to infer the shared state. This state is then mapped through the target cell-type response rule to predict the excluded target response, directly testing whether *W* and *E* generalize to unseen donors. Longitudinal responses are accurately predicted across target cell types in held-out donors (Fig. 7B). When the model is fit with donor correspondence disrupted across cell types within each training visit pair, prediction in correctly matched held-out donors decreases substantially. Thus, longitudinal immune change contains coordinated donor-level information that is shared across cellular populations but related through cell-type-specific response rules; this organization generalizes to unseen individuals.

The recovery of the same separable organization in Sound Life motivates a stronger test of compatibility between *in vivo* and *in vitro* responses. Because the two cohorts span very different distributions of response states, direct training in one setting and transfer to the other conflates compatibility of the cell-type-specific organization with extrapolation across response regimes. We therefore instead ask whether both settings can be described by a single set of cell-type-specific response rules. We use a common PCA basis derived by combining donor-specific trajectories with donor-perturbation responses and fit the cell-type-specific mappings and encoders as before. In donor-level cross-validation, held-out target responses from both cohorts are predicted as before (Fig. 7C, top). In the corresponding donor-permutation control, donor correspondence is disrupted independently across cell types within each training cytokine or longitudinal visit pair before refitting the joint model. Prediction decreases in both cohorts, showing that the shared model depends on coordinated donor-level response structure. Prediction under the permuted control remains higher in Parse because cytokine identity is preserved, retaining stereotyped response structure for which Sound Life has no analogous component.

Having learned common response rules across controlled perturbation and healthy longitudinal variation, we next freeze the joint model and test whether this organization transfers to an independent disease-associated longitudinal cohort spanning progression to rheumatoid arthritis (RA) [33]. For each donor-specific trajectory and target cell type in the RA cohort, the shared state is inferred from the observed non-target cell types and decoded through the frozen target-cell response rule to predict the excluded response (see Eq. 3). The frozen model accurately predicts longitudinal transcriptional changes in the unseen RA cohort and improves prediction over direct cross-cell correspondence (Fig. 7C, bottom). Across 1,188 target-cell predictions spanning 132 donor-specific pairs of visits and nine cell types in the RA cohort, the frozen model improves prediction in 1,161 cases (98%), with strongest gains for monocyte responses. These results show that the jointly learned mappings and encoders transfer beyond both the *in vitro* perturbation and healthy longitudinal settings used for model fitting.

Together, these results show that longitudinal immune change exhibits a separable organization compatible with the structure identified *in vitro*. Controlled perturbation provides a particularly informative setting for resolving cell-type-specific response rules because individual perturbations may isolate specific dimensions of response, making cell-type-specific response differences more apparent. The longitudinal analyses then test whether this organization extends to natural immune change *in vivo*. The transfer of the jointly learned mappings and encoders to the RA cohort further supports a shared organizational framework spanning experimental perturbation, healthy longitudinal variation and longitudinal immune change in a disease setting. Across these contexts, coordinated immune change exhibits a shared-state organization with donor- and perturbation-independent cell-type-specific transcriptional response rules.

### Separable organization predicts unobserved cellular contexts

A direct consequence of the separable organization identified here is that responses need not be measured exhaustively across cellular contexts. If the response rule for a target cell type is stable across donors and perturbations, then observations from a limited set of perturbations can constrain that rule, while responses measured in other observed cell types can be used to infer the shared response state for donor–perturbation combinations in which the target response is unobserved. The target response should then be predictable by combining this inferred state, which is constrained by observed cell types, with the learned target-cell response rule (Fig. 8A). Thus, cross-cell-type context generalization provides an independent test of whether the separable structure captures enough of the response organization to predict cellular contexts that were only sparsely observed during model fitting.

**Figure 8:**
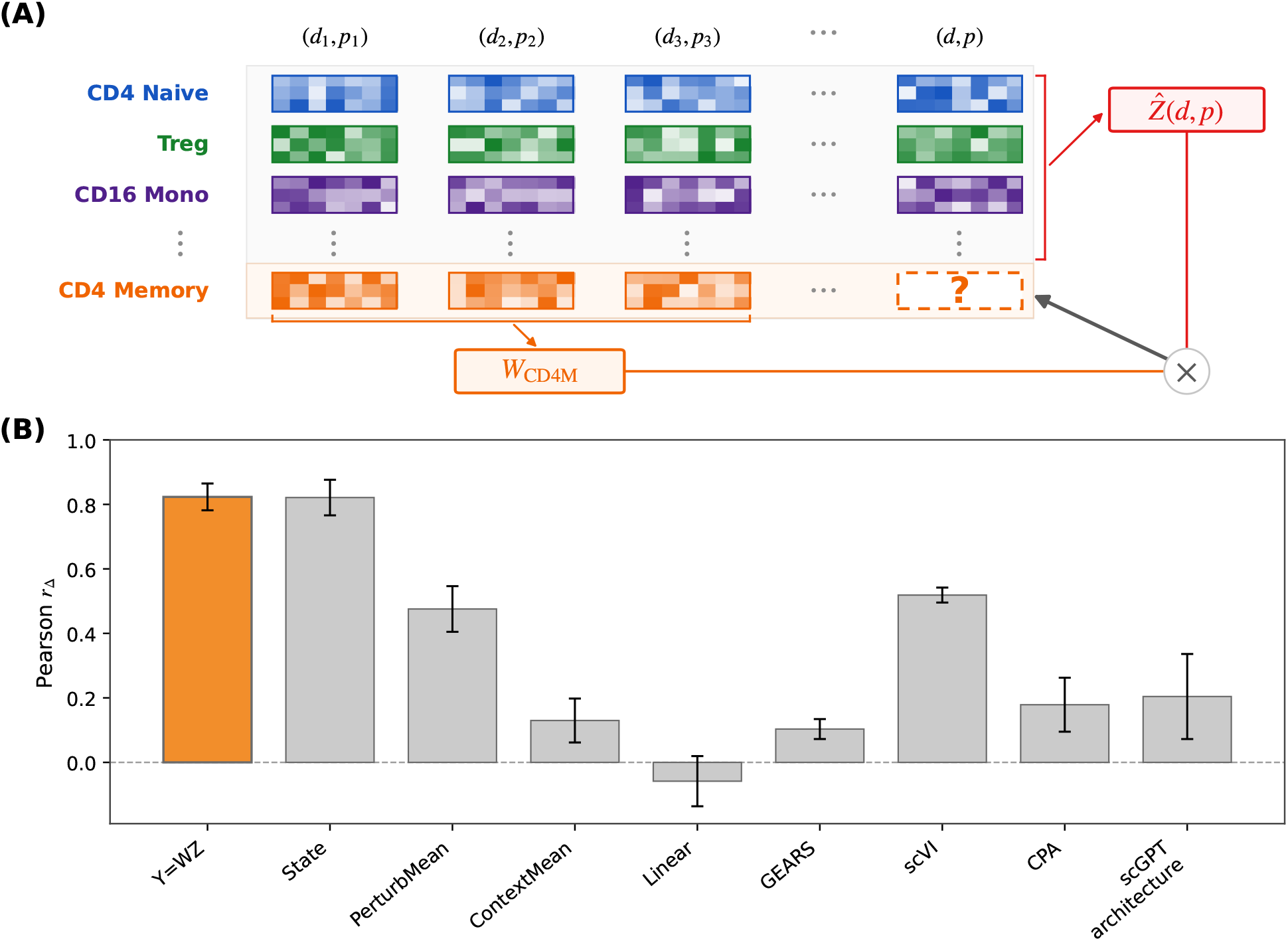
Separable response organization supports prediction of sparsely observed cellular contexts. **(A)** Schematic of target-cell response prediction in the State Parse-PBMC benchmark [39]. For a target cell type, responses to a subset of perturbations constrain its cell-type-specific response rule *W* (*c*), while responses from the remaining observed cell types for a donor–perturbation pair are encoded and averaged to infer the shared response state 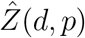. The inferred state is then decoded through the target-cell response rule to predict the missing response. **(B)** Prediction performance on the State Parse-PBMC benchmark. Parse is evaluated using the same target-cell perturbation splits and 2,000 highly variable genes used in the benchmark. Performance is shown together with the models and baselines reported by State. Bars show the mean Pearson correlation between predicted and observed perturbation responses across the four target-cell contexts and error bars show the standard deviation across contexts.

To test cross-cell-type context generalization, we evaluate our model using the Parse-PBMC context-generalization benchmark introduced by State [39]. Because the benchmark and its train/test splits were defined independently of our model, it provides an external test of whether separable organization is sufficient to support context generalization. For each of four target-cell tasks, the target cell type is observed under only 24 training perturbations, with four additional perturbations reserved for validation and responses to 62 perturbations held out for testing^1^, while the remaining cell types retain observations across the full perturbation panel. Using the published splits and 2,000 highly variable genes from State, we fit the model separately for each target-cell task following the benchmark design. We use the same model specification as in our primary Parse analyses, without benchmark-specific hyperparameter tuning or model selection. Our model achieves performance comparable to State on this benchmark (Fig. 8B), showing that the separation of response variation into a shared donor–perturbation state and stable cell-type-specific response rules is sufficient to support strong prediction when perturbation coverage is sparse in the target cell type.

The strong performance on the State benchmark shows that separability is not only descriptive but practically useful for cross-context prediction. Our linear model explicitly represents shared donor-level response variation and cell-type-specific response rules. Its predictive performance can therefore be interpreted in terms of the separable organization encoded by the model. The close agreement in performance between our model and State – a transformer-based perturbation model – suggests that this separable structure may account for much of the predictive signal leveraged by State for context generalization. Thus, the same organization that explains coordinated donor-level response variation also provides a practical basis for predicting unobserved cellular responses.

## Discussion

Transcriptional change across peripheral immune cell types contains a coordinated donor-level component observed across both controlled *in vitro* perturbation and natural longitudinal variation *in vivo*. This organization is well described by a separable model in which distinct cellular responses are represented as cell-type-specific expressions of a shared donor-level response state. Coordinated immune responses therefore do not require different cell types to respond similarly; distinct transcriptional responses may instead represent cell-type-specific expressions of the same shared response state. Controlled perturbations make this organization particularly clear, whereas longitudinal changes likely integrate multiple biological influences and may therefore exhibit stronger direct correspondence across cell types even when compatible cell-type-specific response rules apply. In this context, our previous work has shown that donor-level transcriptional state is coordinated across immune cell types and persists across time, tissues and molecular modalities [17]. In that setting, differences in transcriptional state are strongly aligned across cell types. Here we focus instead on transcriptional change, where responses can differ substantially across cell types, making the cell-type-specific organization of donor-level variation more apparent.

The separable organization we identify suggests that donor-level variation is coordinated across immune cell types and that the mechanisms translating this shared variation into cell-type-specific responses are sufficiently stable across donors. Systemic physiological and environmental influences, together with age, sex, genetic and non-heritable variation [48, 49, 50, 8, 10, 51], shape immune transcriptional state and may also contribute to shared response variation across immune cell types. Conversely, cell-type-specific response rules may reflect lineage-defining regulatory programs that maintain cell identity through signaling, chromatin and transcriptional regulation [52, 53, 54, 55]. Population-scale studies indicate that these core cell-type regulatory programs are broadly conserved across individuals [56, 57]. Together, systemic influences and conserved lineage programs may provide a biological basis for the separable organization we observe, though we emphasize that the underlying molecular mechanisms remain unidentified.

Recent computational models have sought to predict cellular responses across unseen perturbations and biological contexts [35, 36, 37, 39]. However, increasingly large models and pretraining datasets have not consistently improved perturbation prediction over simple baselines, indicating that scale alone is insufficient for generalization [47, 58]. The separable organization identified here provides a complementary basis for generalization across cellular contexts. A response state inferred from observed cell types can be combined with the cell-type-specific response rules to predict an unobserved cellular response. Consistent with this interpretation, the separable model achieves performance comparable to the leading model in the independently defined Parse-PBMC context-generalization benchmark (Fig. 8). Additionally, our results suggest that successful prediction may require biological context beyond perturbation and cell type. In human peripheral immune cells, this context includes donor-specific information because transcriptional change contains a substantial donor-specific component, even as the cell-type-specific response rules appear to be donor-independent. Models that generalize to new individuals may therefore need to leverage a limited set of donor-specific measurements that help constrain their response state. Conversely, generalization across cellular contexts may depend on learning stable response rules that reflect conservation of the mechanisms through which perturbations propagate, rather than similarity of observed expression distributions alone [59].

Several limitations define the scope of our results. First, our analyses are based on average responses within peripheral immune cell types and therefore establish separable organization at the level of donor-specific mean transcriptional change rather than heterogeneity among individual cells. Second, the perturbation analysis includes only 12 donors and is restricted to PBMC responses to cytokine stimulation. Although the recovery of separable organization in Sound Life and the rheumatoid arthritis cohort extends this structure beyond *in vitro* cytokine perturbations, its generality across tissues, disease settings and other classes of perturbation remains uncertain. Third, it remains unclear what information is necessary and sufficient to constrain donor-specific response state, including how much information baseline donor state provides. Large multi-omic studies combining baseline characterization, longitudinal sampling and controlled *in vitro* perturbation in the same individuals could test the generality of cell-type-specific response rules and identify the molecular, physiological and functional measurements that constrain donor-specific response state. Higher signal-to-noise single-cell measurements could further test whether separable organization extends to variation among individual cells.

We find that transcriptional responses across peripheral immune cell types are coordinated at the donor level and can be represented by a shared response state expressed through cell-type-specific response rules. This organizing principle provides a tractable link across biological scales, showing how distinct cellular responses can reflect coordinated variation at the level of the individual. More broadly, these findings raise the possibility that multicellular systems exhibit a common organizing structure in which cellular populations across tissues and molecular modalities are constrained by shared system-level variation. Understanding this organization may ultimately improve our ability to predict responses to interventions and perhaps reveal new therapeutic targets for reshaping coordinated cellular behavior in disease.

## Materials & Methods

### Datasets

#### Parse cytokine perturbation dataset

We analyze a single-cell peripheral blood mononuclear cell (PBMC) perturbation dataset comprising approximately 10 million cells from 12 donors exposed *in vitro* to 90 cytokines and a phosphate-buffered saline (PBS) control [32]. The same donors are measured across the perturbation panel, providing matched responses across immune cell types within each donor–cytokine combination. Donor–cytokine–cell-type groups containing fewer than 100 cells are excluded. We restrict the analysis to 11 broadly represented immune populations represented in *>* 50% of possible donor-cytokine groups: naive and memory CD4 T cells, naive and memory CD8 T cells, Tregs, MAIT, naive and intermediate/memory B cells, natural killer cells and CD14 and CD16 monocytes.

Expression is initially aggregated across 40,352 measured genes. Gene prevalence is defined across all retained donor–cytokine–cell-type mean profiles in the full Parse dataset. We retain genes with nonzero mean expression in at least 50% of these profiles, yielding 18,992 genes. Restricting the prevalence calculation to the 11 primary populations yields a nearly identical gene set.

#### Sound Life longitudinal dataset

We use an independent longitudinal PBMC cohort from the Sound Life Project to determine whether the organization identified under controlled cytokine stimulation extends to natural immune variation over time [16, 17]. The analyzed cohort contains 95 donors sampled across nine scheduled visits spanning two influenza-vaccination studies and an immune-variation study. The retained visits are Flu Year 1 Days 0, 7 and 90, Flu Year 2 Days 0, 7 and 90 and Immune Variation Days 0, 7 and 90.

Cell identities are defined using the provided Allen Institute for Immunology (AIFI) annotations. We analyze nine populations shared with the Parse perturbation dataset: naive and memory CD4 T cells, naive and memory CD8 T cells, naive and memory B cells, CD56dim natural killer cells and CD14 and CD16 monocytes. For each donor–visit–cell-type group, cells are library-size normalized to 4,524 counts per cell, log transformed and averaged. Groups containing fewer than 100 cells are excluded, yielding 6,984 donor–visit–cell-type expression profiles across the nine populations.

Expression is initially retained across all 33,538 measured genes. We then restrict to genes with nonzero mean expression in at least 50% of the 6,984 aggregated profiles, yielding 15,493 Sound Life genes. For analyses jointly modeling Parse and Sound Life, this set is intersected with the 18,992-gene Parse universe, yielding 12,249 genes shared between the two cohorts.

#### Rheumatoid arthritis longitudinal dataset

We use an independent longitudinal PBMC cohort spanning progression to rheumatoid arthritis (RA) as an external test of response-rule transfer [33]. We analyze the 15 individuals classified as converters, comprising 66 longitudinal samples. Cell identities are defined using the AIFI annotations provided with the study and matched to the raw single-cell count data. Group-level expression profiles are constructed for nine populations shared with the Parse and Sound Life analyses: naive and memory CD4 T cells, naive and memory CD8 T cells, naive and memory B cells, CD56dim natural killer cells and CD14 and CD16 monocytes. Because the RA cohort is smaller and is used only for external evaluation rather than model fitting, we use a lower minimum of 25 cells per donor–visit–cell-type mean than in the Parse and Sound Life analyses. Longitudinal responses are constructed from all within-donor pairs of available visits, yielding 132 donor–visit-pair states across 15 donors. The RA dataset contains 33,538 measured gene features.

### Construction of donor-level perturbation responses

For each Parse cell, transcript counts are divided by the empirical library size, scaled to a total of 4,524 counts and transformed as log(1 + *x*). Normalized expression values are averaged within each donor–cytokine–cell-type group. Groups containing fewer than 100 cells are excluded.

For donor *d*, cell type *c* and cytokine *p*, the perturbation response is defined as

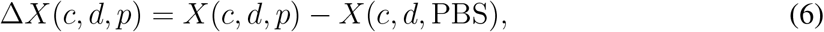

where *X*(*c, d, p*) is the normalized mean expression profile of the stimulated group and *X*(*c, d*, PBS) is the donor- and cell-type-matched PBS profile. The resulting response vectors describe donor- and cell-type-specific transcriptional change relative to the corresponding unstimulated condition.

### Principal-component representation of perturbation responses

Principal-component analysis (PCA) is applied to the donor-level perturbation-response matrix. PCA is fit directly to the gene-space response vectors using randomized singular-value decomposition with random seed 0. The first 300 principal components are retained for the primary Parse analyses. We denote the resulting *K*-dimensional response principal-component representation by *Y* (*c, d, p*). PCA performs its standard feature-wise mean centering, but the resulting principal-component coordinates are not subsequently standardized or rescaled. The scale of each component therefore retains information about the amount of response variance represented along that axis.

The main conclusions are evaluated for sensitivity to the precise number of retained components. For descriptive full-data analyses, PCA is fit using all available perturbation responses. For joint donor–perturbation analyses in Figures 2 and 3, PCA is refit within each training split and held-out responses are projected using the training-derived basis.

### Direct cross-cell-type coordination

Direct coordination is measured using Pearson correlation across the first 300 response principal components. Correlations are computed separately for individual donor–cytokine response vectors using the original, unstandardized principal-component coordinates.

For the cell-type similarity matrix, each cell-type pair is evaluated using all donor–cytokine combinations in which both cell types are observed; observations are not required to contain the other analyzed cell types. For each matched donor–cytokine observation, Pearson correlation is computed between the corresponding response vectors. Each matrix entry is the median correlation for that cell-type pair across available donor–cytokine observations.

To distinguish donor-specific coordination from the stereotyped response to each cytokine, three comparisons are made. First, for each cytokine and donor, within-donor coordination is calculated across all available cell-type pairs and summarized by the median, followed by the median across donors for each cytokine. Second, donor correspondence is broken while preserving cytokine identity and cell-type-pair weighting. For each donor and available cell-type pair, one member of the pair is replaced by the corresponding response from each eligible different donor exposed to the same cytokine, considering both pair orientations. These correlations are summarized first across unmatched donors for each cell-type pair, then across cell-type pairs within each donor and finally across donors for each cytokine. Third, a global unmatched reference is generated from 50,000 random cross-cell-type pairs for which both donor identity and cytokine identity differ. Random sampling uses seed 0.

### joint donor–perturbation cross-validation

Generalization across donors and perturbations is evaluated using a common joint donor–perturbation cross-validation scheme for the analyses in Figures 2 and 3. Cytokines are shuffled with seed 0 and divided into 10 folds. Within each perturbation fold, each of the 12 donors is held out in turn, yielding 120 training–test splits. For each split, the training data contain responses from the other 11 donors under the 81 training cytokines, while evaluation is performed on responses from the held-out donor under the nine held-out cytokines. The same donor and perturbation splits are used for the cross-cell-type mapping and shared-state analyses.

Any response representation learned within these analyses is fit using only the corresponding training data. In particular, the response PCA is refit independently within each split and held-out responses are projected into the training-derived response-PC basis.

### Cross-cell-type response mappings

To test whether one cellular response can be translated into another, we fit an affine ridge-regression mapping from a source cell type *c*_*s*_ to a target cell type *c*_*t*_:

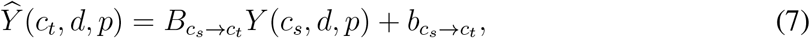

where *Y* (*c, d, p*) ∈ ℝ^*K*^ denotes the response principal-component vector. Ridge regression is fit with an *L*_2_ penalty of 1 and an intercept. Principal-component coordinates are not standardized before fitting.

Generalization across donors and perturbations is assessed using the joint donor–perturbation cross-validation scheme defined above. For each split, a mapping is fit using paired donor– cytokine responses from the 11 training donors and 81 training cytokines and applied without modification to responses from the held-out donor under the nine held-out cytokines. Prediction accuracy is calculated separately for each held-out donor–cytokine response as the Pearson correlation across principal components between the predicted and observed target responses. For the CD4 memory-to-CD14 monocyte example, mapped prediction is compared with direct, untransformed correspondence between the observed source and target responses. These values are summarized by their median across held-out donors for each cytokine. For the all-cell-type analysis, prediction correlations are pooled across held-out donor–cytokine observations and summarized by the median for each source–target pair.

### Donor-specific response variation

To isolate donor-specific variation beyond the stereotyped cytokine response, the mean response to each cytokine is estimated separately within each cell type from the other donors exposed to the same cytokine. The superscript (−*d*) denotes that the held-out donor *d* is excluded from this estimate:

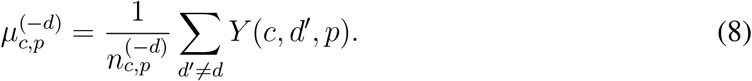

For held-out donor *d*, the donor-specific response deviation is defined as

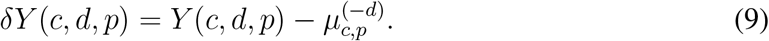

For the joint donor–perturbation analysis, responses from other donors under the held-out cytokine are used only to estimate the stereotyped cytokine response and define the donor-specific deviations; they do not contribute to fitting the response PCA or cross-cell-type mapping.

The affine source–target mapping is fit to the full response vectors *Y* (*c, d, p*) from the corresponding joint donor–perturbation training split. Its linear component is then applied without refitting to the held-out donor-specific deviation:

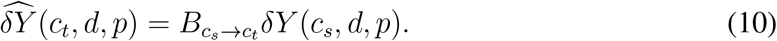

Direct correspondence between source and target deviations is compared with correspondence after applying the learned mapping. Correlations are calculated separately for each held-out donor–cytokine observation and summarized by the median for each source–target pair.

Dependence on donor correspondence is evaluated by comparing each mapped donor-specific response deviation with the target deviation from the matched donor and with deviations from other donors exposed to the same cytokine. For each prediction, the unmatched-donor reference is the median Pearson correlation across all available target deviations from different donors with the same cytokine. Matched and unmatched prediction accuracy are then summarized separately for each source–target cell-type mapping.

### Shared donor–perturbation state model

For donor *d*, perturbation *p* and cell type *c*, let *Y* (*c, d, p*) ∈ ℝ^*K*^ denote the observed response in response-PC space, with *K* = 300. We model the responses as

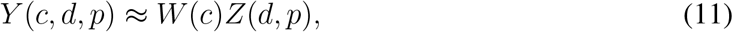

where *Z*(*d, p*) ∈ ℝ^*q*^ is a donor–perturbation response state shared across cell types and *W* (*c*) ∈ ℝ^*K×q*^ is a cell-type-specific response rule. We use *q* = 300 for the primary analysis. Thus, the model does not impose dimensional reduction within an individual cellular view; rather, it constrains the 11 *K*-dimensional cell-type-specific responses for each donor–perturbation pair to be represented by a single *K*-dimensional shared state and cell-type-specific response rules.

The model is fit using uncertainty-weighted alternating least squares. Let *ω*_*c,d,p,k*_ denote the weight assigned to response component *k* for cell type *c*, donor *d* and perturbation *p*. The fitted objective is

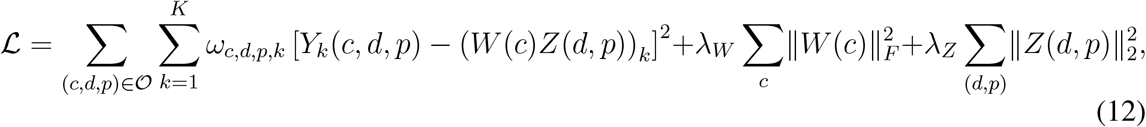

where *O* denotes the set of observed donor–perturbation–cell-type combinations. Missing cell-type views do not contribute to the objective.

Gene-level uncertainty in each donor–cytokine–cell-type response is estimated from 1,001 bootstrap resamplings of the contributing cells. The resulting response variances are propagated into response-PC space using the squared PCA loadings:

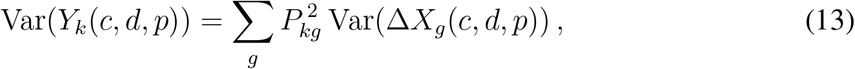

where *P*_*kg*_ is the loading of gene *g* on response component *k*. These variances are used to define the inverse-variance weights *ω*_*c,d,p,k*_ in the model objective.

The model is initialized with random Gaussian response-state coordinates using seed 0. Alternating updates solve weighted ridge-regression problems for *W* (*c*) conditional on *Z*(*d, p*) and for *Z*(*d, p*) conditional on *W* (*c*). For the Parse analyses, we use *λ*_*W*_ = *λ*_*Z*_ = 1 and five alternating iterations. Response PCs, cell-type views and shared-state coordinates are not standardized.

### Inference of shared state from observed cell types

For each cell type, we fit a separate multi-output ridge encoder *E*(*c*) that maps its observed response into the fitted shared-state space:

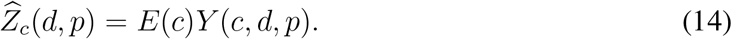

Each encoder maps the 300 response PCs of one cell type to the fitted 300-dimensional shared state and is trained with an *L*_2_ penalty of 1 and without an intercept. Across the shared-state analyses, the decoder, state and encoder regularization parameters are fixed a priori at 1 and are not tuned using cross-validation or held-out prediction performance.

For prediction of a target cell type *c*_*t*_, the shared state is inferred by averaging the encoded state estimates from the other observed cell types:

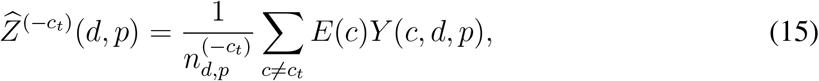

where the sum is over observed cell types for donor–perturbation pair 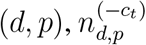 is the number of observed non-target cell types and the superscript (−*c*_*t*_) denotes that the target cell type *c*_*t*_ is excluded from the estimate. The target response is then predicted as

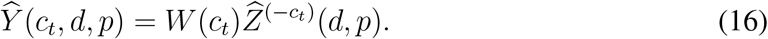

The target cell type is again excluded from state inference in all held-out multiview prediction analyses.

### Model reconstruction and joint donor–perturbation transfer

Reconstruction accuracy is evaluated for each observed donor–perturbation–cell-type response by comparing *Y* (*c, d, p*) with *W* (*c*)*Z*(*d, p*) using Pearson correlation across the 300 response PCs. Generalization across donors and perturbations is evaluated using the joint donor–perturbation cross-validation scheme defined above. For each split, the shared-state model, cell-type response rules and encoders are learned using only responses from the corresponding 11 training donors and 81 training cytokines.

Evaluation is restricted to responses from the held-out donor under the held-out cytokines. For each held-out donor–cytokine pair and target cell type, prediction is performed using the target-held-out multiview inference procedure defined above. The shared-state model, target decoder and source encoders are therefore learned without either the test donor or the test cytokine. Accuracy is calculated separately for each held-out donor–cytokine–target-cell-type response as the Pearson correlation across response PCs between predicted and observed target responses and is summarized by target cell type.

A donor-correspondence control is constructed independently for every donor–perturbation cross-validation split. After restricting the data to the same 11 training donors and 81 training cytokines used for the corresponding observed fit, donor assignments are independently deranged within each cytokine–cell-type stratum. This preserves the response vectors, cytokine identity, cell-type-specific response distributions, donor composition and observation counts within each training stratum while disrupting donor correspondence across cellular views. The complete response representation, shared-state model, cell-type response rules and encoders are then refit on these donor-mismatched training responses and evaluated on the same correctly matched held-out donor–cytokine responses using the identical target-held-out multiview inference procedure. Derangements are generated reproducibly using fixed random seeds.

### Response-variance decomposition and measurement uncertainty

To quantify how much of the observed transcriptional response variance is reproducible and how much of that reproducible variation is captured by the separable model, we construct independent split-half response measurements from the underlying single-cell data [45]. For each donor, perturbation and cell type, cells are randomly partitioned into two approximately equal subsets, denoted *A* and *B*. Stimulated and matched PBS cells are split independently, with the same PBS split reused across cytokines for a given donor and cell type within each replicate split. Each half is normalized and log-transformed using the same procedure as the primary response data, and the corresponding stimulated-minus-PBS mean response is calculated. Five independent *A/B* split pairs are generated using fixed random seeds.

Let 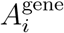 and 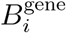 denote the two gene-space half-sample responses for donor–perturbation– cell-type response *i*. If the full-sample response is written as a reproducible component *S*_*i*_ plus measurement error, the two half-sample responses can be written approximately as

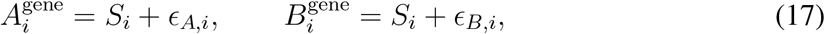

where the two measurement errors are independent between split halves. Because each half contains approximately half of the available cells, the variance of each half-sample mean is approximately twice that of the corresponding full-sample mean. The measurement variance of the full-sample response can therefore be estimated directly from split-half disagreement as

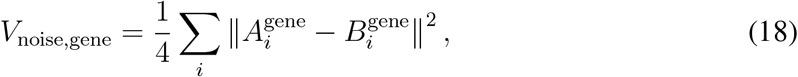

where *i* indexes donor–perturbation–cell-type responses. Reproducible gene-space response variation is estimated using the corresponding cross-replicate product,

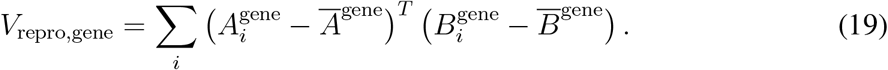

The two split-half response matrices are centered separately across responses before calculating the cross-replicate product. These quantities are calculated separately for each of the five independent split pairs and averaged across splits. As an independent check, measurement uncertainty is also calculated analytically from the empirical cell-level covariance and the number of stimulated and PBS cells contributing to each mean. The split-half and analytic estimates agree closely.

All response-variance quantities are expressed relative to the total observed gene-space response variance. Let *X*_*i*_ denote the full-sample response vector and 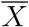 the mean response across all observed donor–perturbation–cell-type samples. We define

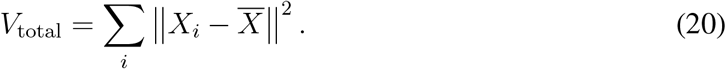

The fraction of total response variance attributable to measurement uncertainty is therefore *V*_noise,gene_*/V*_total_, with the remaining variance treated as reproducible response variation.

Because principal components are selected to maximize variance in the same response matrix used for evaluation, using the full-data PCA basis to partition reproducible variation inside and outside the retained response space can preferentially retain realized sampling fluctuations.

We therefore estimate this partition using leave-one-donor-out cross-fitted PCA. For each donor *d*, a response PCA is fit using responses from the other 11 donors, with standard feature-wise mean centering and no gene scaling. The independently generated *A* and *B* responses from donor *d* are then projected into this training-donor PCA basis. Thus, the basis used to evaluate donor *d* is independent of both the donor’s biological response variation and its realized measurement error.

Let 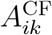 and 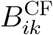 denote the coordinates of the two half-sample responses along cross-fitted response component *k*. For a retained dimensionality *K*, reproducible response variation represented within the PCA space is estimated as

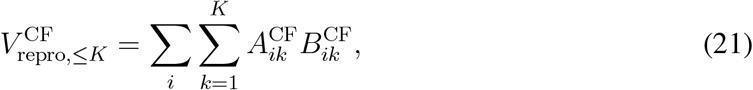

and the fraction of all reproducible response represented within *K* dimensions is

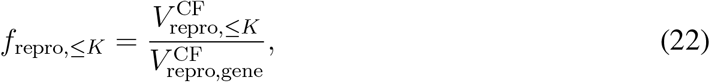

where 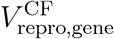 denotes the corresponding cross-fitted estimate of total reproducible gene-space response variation. The fraction outside the retained space is 1 − *f*_repro,≤*K*_. Measurement uncertainty is partitioned analogously using

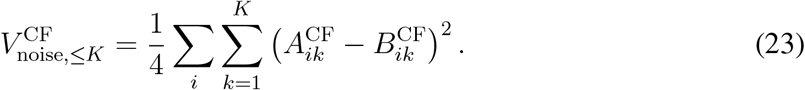

The cross-fitted fractions of reproducible signal and measurement uncertainty lying inside and outside the retained space are then mapped onto the total response-variance scale defined above. At *K* = 300, this procedure indicates that 94.7% of reproducible response variation lies within the retained response space, whereas only 9.6% of estimated total gene-space measurement uncertainty lies within this space.

For each retained response dimensionality *K*, the shared-state model is fit with *q* = *K* in the fixed response-PC basis defined by the full-data PCA, truncated to its first *K* components.

For each held-out donor, the shared-state model and encoders are fit using responses from the remaining 11 donors, with all perturbations retained. We then compare the shared-state model with two nested leave-one-donor-out stereotype predictors to quantify how much reproducible variation it captures beyond increasingly specific stereotyped responses. For held-out donor *d*, cytokine *p* and cell type *c*, the cell-type-specific stereotyped response is defined as

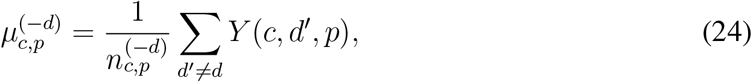

where the superscript (−*d*) denotes that the held-out donor *d* is excluded from this estimate. A cytokine-only stereotyped response is then obtained by averaging the cell-type-specific means equally across the 11 cell types,

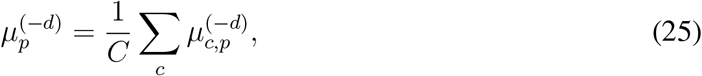

with *C* = 11. Equal weighting prevents small differences in the number of observed donor– cell-type combinations from changing the definition of the generic cytokine response.

For these calculations, let 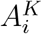 and 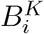 denote the two split-half responses projected into the fixed *K*-dimensional response-PC basis used for the shared-state model, and let 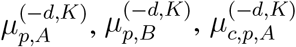 and 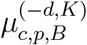 denote the corresponding split-specific stereotype predictors. Reproducible variation within this fixed response-PC space is

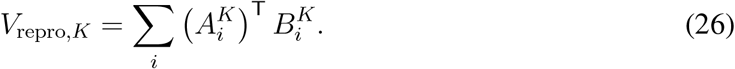

For each split, we calculate the reproducible residual variation remaining after the cytokine-only predictor,

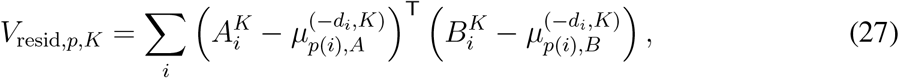

and after the cell-type-specific cytokine predictor,

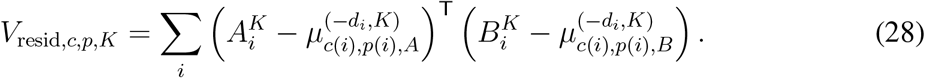

The full *WZ* predictor is evaluated using these leave-one-donor-out models. For each heldout donor–perturbation pair and target cell type, the target response is excluded from state inference, the shared state is estimated from the available non-target cell types and the target response is generated through the corresponding decoder. Applying the saved fold-specific model separately to 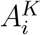 and 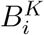 gives predictions 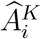 and 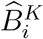, from which the reproducible model residual is

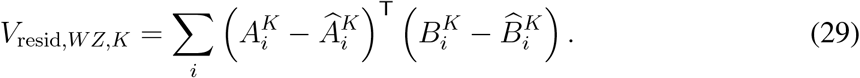

We use differences among these residual quantities to define nested increments in reproducible response variance. Cytokine stereotypy is

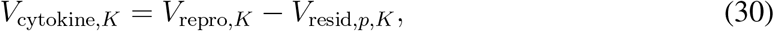

the additional contribution of stereotyped cell-type-specific response is

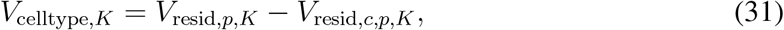

and the additional reproducible response captured by the shared-state model beyond the leave-one-donor-out cell type–cytokine mean is

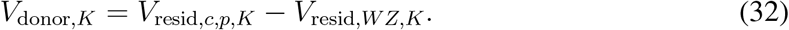

The sum

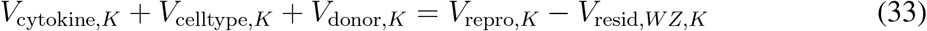

is therefore the total reproducible response variance captured by the *WZ* predictor within the retained response space. These quantities are nested increments in predictive variance rather than an orthogonal ANOVA decomposition and are interpreted accordingly.

The *WZ* prediction analysis is performed in the fixed response-PC basis used for the primary model, whereas the fraction of reproducible variation represented within each *K* is estimated using donor-cross-fitted PCA to avoid PCA-selection bias. To place both quantities on a common response-variance scale, we preserve the relative decomposition of reproducible variance within the fixed *K*-dimensional model space and rescale its absolute magnitude to the cross-fitted estimate of reproducible variation represented within the same *K*. Reproducible response outside the retained space and measurement uncertainty inside and outside the retained space are then added to obtain a decomposition that closes to the total observed response variance. Because this final rescaling combines model performance evaluated in the fixed PCA basis with representation fractions estimated from cross-fitted PCA, the resulting fraction of all reproducible variance captured by *WZ* is treated as an estimate. In contrast, the fraction of reproducible variance captured within the represented *K*-dimensional response space is evaluated directly from the leave-one-donor-out prediction residuals.

### Interpretation of cell-type response rules and shared states

Because the factorization does not assign unique biological meaning to individual latent coordinates, interpretation focuses on relationships among fitted cell-type response rules, gauge-invariant source-to-target operators and relative organization within the shared-state space. The full-data model used for these analyses is fit using all donors and cytokines with *K* = *q* = 300, *λ*_*W*_ = *λ*_*Z*_ = 1 and the same weighted alternating least-squares procedure described above.

Similarity between cell-type response rules is calculated by flattening each fitted *W* (*c*), normalizing it to unit Frobenius norm and computing pairwise cosine similarity.

To characterize the observable mapping between cell types, we combine the fitted source encoder with the target response rule. In column-vector notation, the resulting source-to-target operator is

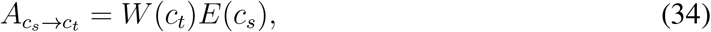

such that a response observed in source cell type *c*_*s*_ is mapped to the corresponding response in target cell type *c*_*t*_. Although *W* (*c*), *E*(*c*) and *Z*(*d, p*) depend on the choice of latent coordinates, the composed operator 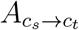 is invariant to a common orthogonal reparameterization of the latent state and therefore defines a direct mapping in the observed response space.

For visualization, we examine the mappings between CD4 memory T-cell and CD14 monocyte responses in both directions. The signed coefficients of the leading 15 *×* 15 response-PC blocks of 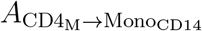 and 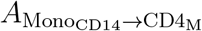 are displayed without normalization and with a shared coefficient scale. Rows denote target response principal components and columns denote source response principal components. The full operators are 300 *×* 300; only the leading response-PC blocks are shown for visualization.

To isolate perturbation-associated structure in *Z*(*d, p*), we first remove each donor’s mean response state across perturbations. The perturbation effect for cytokine *p* is then

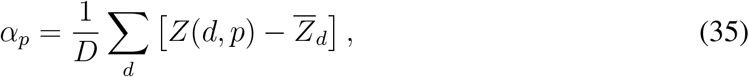

where

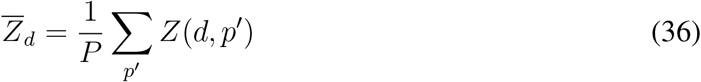

is the mean response state for donor *d* across perturbations. Because all 12 donors are represented for all 90 perturbations, this is equivalent to subtracting the mean state across all donor–perturbation pairs from the donor-averaged state for each cytokine. Cytokine effects are normalized to unit length and compared using cosine similarity. Cytokines are ordered by average-linkage hierarchical clustering of cosine distance and annotated using the perturbation groups defined in the source study.

To examine donor-associated structure, the mean state for each cytokine is removed:

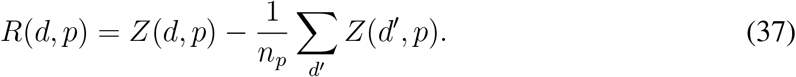

Residual states are normalized to unit length and compared using cosine similarity. States are ordered by donor and then cytokine. Donor organization is summarized by comparing within-donor and between-donor cosine similarities and by determining whether the nearest neighboring residual state belongs to the same donor.

### Construction of longitudinal immune changes

All 36 unordered pairs of the nine visits are considered. For visits *v*_*a*_ and *v*_*b*_, with *v*_*b*_ later in the predefined visit ordering, longitudinal change is defined as

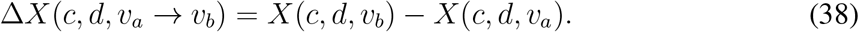

For each donor and visit pair, the collection of cell-type-specific longitudinal changes defines a donor-specific trajectory. A cell-type-specific change is retained whenever both visits are available for that donor and cell type; availability of the other cell types is not required. The sign convention is applied consistently across cell types. Because each donor contributes multiple overlapping visit pairs, these donor-specific trajectories are treated as pairwise longitudinal differences rather than statistically independent biological trajectories.

### Direct coordination of longitudinal change

For the descriptive coordination analysis, we use all available cell-type-specific responses from the Sound Life donor-specific trajectories. PCA is fit to the 12,249-gene longitudinal-response matrix using randomized singular-value decomposition with random seed 0, and the first 300 components are retained. We denote the resulting response-PC representation by *Y* (*c, d*, Δ*t*), where Δ*t* denotes the visit pair defining the donor-specific trajectory. The resulting principal-component coordinates are not standardized.

For each cell-type pair, we retain all donor-specific trajectories in which both cell types are observed; trajectories are not required to contain the other analyzed cell types. Pearson correlation is computed across the 300 response PCs separately within each trajectory. Each cell-type matrix entry is the median correlation for that pair across all available trajectories.

### Strict donor-level cross-validation in Sound Life

Generalization to unseen donors is assessed using five-fold cross-validation. Donors, rather than individual visit-pair observations, are divided into folds using shuffled K-fold splitting with seed 0. All observations from each held-out donor are assigned to the same fold.

Within each fold, PCA is fit using only longitudinal-difference vectors from training donors and held-out responses are projected into the training-derived basis. A shared-state model with *K* = *q* = 300 is then fit using only training donors. The Sound Life model uses five alternating iterations, *λ*_*W*_ = *λ*_*Z*_ = 1 and ridge encoders with penalty 1. No response-PC, cell-type-view or shared-state standardization is applied. Uniform uncertainty values are used, making the reconstruction term effectively equally weighted.

For each held-out donor-specific trajectory and target cell type *c*_*t*_, prediction is performed using the target-held-out multiview inference procedure defined above (Eqs. 15–16), with Δ*t* denoting the visit pair defining the trajectory. The target response is excluded from state inference, the shared state is inferred from the available non-target cell types and the resulting state is decoded through *W* (*c*_*t*_) to predict the excluded target response. Prediction accuracy is calculated separately for each held-out donor–visit-pair–target-cell-type response as the Pearson correlation across response PCs between predicted and observed target responses. Out-of-fold correlations are pooled across the five folds and summarized separately for each target cell type.

### Sound Life donor-correspondence control

Donor correspondence is tested by disrupting cross-cell donor alignment during model fitting. Within each cross-validation fold and exact visit pair, donor responses are independently deranged within each cell type so that the training views assigned to a nominal donor are assembled from responses originating from different donors. This preserves the donor holdout structure, visit-pair identity, cell-type-specific response distributions, the training-only PCA basis and the number of training observations while removing coordinated donor correspondence across cell types.

A single reproducible donor-shuffle realization is generated using seed 10000. The shared-state model and cell-type encoders are refit on the donor-mismatched training data and evaluated on the original correctly matched held-out donors. Evaluation is identical to that used for the unshuffled model: for each target cell type, the shared state is inferred by averaging encoded states from all observed non-target cell types and is then decoded into the excluded target. Prediction accuracy is calculated for each held-out donor–visit-pair combination as Pearson correlation across response PCs and pooled across cross-validation folds.

### Joint Parse–Sound Life model

Compatibility between the two response settings is tested by fitting a single shared-state model jointly to Parse cytokine responses and Sound Life longitudinal differences. The analysis is restricted to nine matched cell types and genes shared between the cohort-specific gene sets.

Generalization is assessed using 12-fold donor-level cross-validation. Each fold holds out one Parse donor and a disjoint subset of Sound Life donors, with every donor held out exactly once. Within each fold, a common PCA basis with *K* = 300 components is fit jointly using training responses from both cohorts. A single shared-state model with *q* = 300 is then fit to the combined training data using one common set of cell-type-specific response rules and one encoder per cell type. The decoder and state regularization parameters are set to *λ*_*W*_ = *λ*_*Z*_ = 1, the encoder ridge penalty is set to 1 and five alternating iterations are used. Uniform uncertainty values are used for both cohorts, making the reconstruction term effectively equally weighted across response components. Responses from the two cohorts are concatenated for model fitting, with each observed cell-type-specific response from either cohort contributing equally to the reconstruction objective; no additional weighting is applied to balance the cohorts.

Held-out donor–perturbation pairs in Parse and held-out donor-specific trajectories in Sound Life are evaluated separately using the target-held-out multiview inference procedure defined above (Eqs. 15–16). Prediction accuracy is calculated separately for each held-out target-cell response as the Pearson correlation across response PCs between predicted and observed responses. Out-of-fold predictions are pooled within each cohort. As a donor-correspondence null, donor assignments are independently deranged across cell types within each cohort and exact training condition, as above: within cytokine for Parse and within visit pair for Sound Life. The training-derived response PCA basis is held fixed, while the joint shared-state model and cell-type encoders are refit on the donor-mismatched training responses and evaluated on the same correctly matched held-out data.

### External transfer to rheumatoid arthritis progression

To test whether the response organization learned from Parse and Sound Life generalizes to an independent biological setting, we fit a final shared-state model to the full Parse and Sound Life datasets and apply it without refitting to longitudinal responses from the rheumatoid arthritis progression cohort [33]. The analysis uses the nine cell types shared among Parse, Sound Life and RA, a common response-PC basis with *K* = 300, a shared-state dimension of *q* = 300, *λ*_*W*_ = *λ*_*Z*_ = 1, an encoder ridge penalty of 1 and five alternating iterations. Uniform uncertainty values are used for both Parse and Sound Life when fitting this joint model.

External transfer begins from the 12,249-gene Parse–Sound Life universe. Genes are matched to the RA data by gene symbol; symbols that are absent from RA or map ambiguously to multiple RA gene features are excluded from all three datasets before model fitting. The response PCA basis, cell-type-specific response rules and cell-type encoders are fit using only the combined Parse and Sound Life responses and are then held fixed for all RA analyses.

For each RA donor-specific trajectory and target cell type, prediction is performed using the frozen target-held-out multiview inference procedure defined above (Eqs. 15–16). All response representations, encoders and cell-type response rules are learned from Parse and Sound Life and held fixed during RA evaluation. Prediction accuracy is quantified as the Pearson correlation across response PCs between predicted and observed target responses. As a direct cross-cell baseline, the eight non-target response vectors are averaged in the same frozen response-PC space and compared directly with the observed target response. This baseline tests prediction from shared response similarity without applying the learned cell-type-specific response rules. Robustness to cell-count variation is assessed by repeating the evaluation after requiring progressively larger minimum cell counts at both endpoints of every longitudinal comparison.

### State Parse-PBMC benchmark

To evaluate the separable model on an independently defined context-generalization task, we reproduce the Parse-PBMC benchmark from State [39]. The benchmark defines four target-cell tasks: B intermediate/memory, B naive, CD4 memory and CD14 monocytes. Each target cell type is evaluated in a separate model fit using the published perturbation partition of 24 training, four validation and 62 test perturbations. For a given target-cell task, responses to the validation and test perturbations are excluded only for the target cell type; responses from the remaining Parse cell types under those same perturbations remain available. The analysis is restricted to the 11 canonical Parse populations and the 2,000 highly variable genes used in the State benchmark.

For each target-cell task, the response PCA, shared-state model and encoders are refit using the corresponding training data and the same model specification described above. Held-out target responses are predicted using the target-held-out multiview inference procedure in Eqs. 15–16. To match the State evaluation, predicted donor-level perturbation responses are transformed back to the 2,000-gene expression space and added to the corresponding donor- and cell-type-matched PBS expression. Predicted and observed stimulated profiles are pooled across donors using stimulated-cell counts, while PBS profiles are pooled using PBS-cell counts. Prediction accuracy for each held-out perturbation is the Pearson correlation across genes between the predicted and observed signed perturbation responses after subtraction of the pooled PBS profile. Correlations are averaged across the 62 test perturbations within each target-cell task. Figure 8 reports the mean and standard deviation across the four target-cell tasks. Performance values for State and the other comparison methods are extracted from the published State Figure 2E.

### Evaluation metrics and statistical summaries

Pearson correlation is the primary evaluation metric for response reconstruction and prediction. Unless otherwise stated, correlation is computed separately for each donor-level response vector across retained principal components. Multiple independent donor observations are not flattened into a single correlation. Distributions are summarized using medians and interquartile ranges.

Cosine similarity is used to compare fitted response rules, source-to-target operators and shared-state vectors where orientation rather than component-wise centering is the quantity of interest. Mean-squared error and *R*^2^ are calculated as secondary diagnostics. For joint donor– perturbation prediction in Figure 3, we report a pooled zero-baseline *R*^2^,

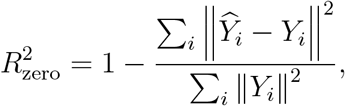

where *i* indexes held-out donor–cytokine–cell-type responses. This statistic measures the fraction of held-out response variance captured relative to the zero vector in the training-derived response-PC space.

Permutation controls are used to determine how much correspondence remains after donor, cytokine or visit-pair matching is disrupted. These controls are interpreted as empirical reference distributions rather than formal tests unless an empirical *p* value is explicitly reported.

### Software and reproducibility

Analyses are performed in Python using NumPy, pandas, SciPy and scikit-learn. Random seeds are fixed for PCA, cross-validation, model initialization and permutation analyses.

## Data Availability

All data used in this study are publicly available. Details of the Parse Biosciences, Sound Life cohort and Rheumatoid Arthritis cohort data may be found in Oesinghaus et al. [32], Gong et al. [16] and He et al. [33], respectively.

## Declaration of generative AI and AI-assisted technologies in the manuscript preparation process

During the preparation of this work the author used ChatGPT 5.6 (OpenAI) in order to generate code and edit manuscript for clarity. After using this tool, the author reviewed and edited the content as needed and takes full responsibility for the content of the published article.

## Acknowledgments

We thank Miah Wander for comments and discussion that improved the manuscript.

## Footnotes

1 The four validation perturbations are not used by our model for either training or testing.

